# Transcriptome profiling reveals CD73 and age-driven changes in neutrophil responses against *Streptococcus pneumoniae*

**DOI:** 10.1101/2021.04.14.439887

**Authors:** Manmeet Bhalla, Lauren R. Heinzinger, Olanrewaju B. Morenikeji, Brandon Marzullo, Bolaji N. Thomas, Elsa N. Bou Ghanem

## Abstract

Neutrophils are required for host resistance against *Streptococcus pneumoniae* but their function declines with age. We previously found that CD73, an enzyme required for antimicrobial activity, is down-regulated in neutrophils from aged mice. This study explored transcriptional changes in neutrophils induced by *S. pneumoniae* to identify pathways controlled by CD73 and dysregulated with age. Ultrapure bone marrow-derived neutrophils isolated from wild type (WT) young, old, and CD73KO young mice were mock-challenged or infected with *S. pneumoniae ex vivo*. RNA sequencing was performed to identify differentially expressed genes (DEGs). We found that infection triggered distinct global transcriptional changes across hosts, that were strongest in CD73KO neutrophils. Surprisingly, there were more down-regulated than up-regulated genes in all groups upon infection. Down-regulated DEGs indicated a dampening of immune responses in old and CD73KO hosts. Further analysis revealed that CD73KO neutrophils expressed higher numbers of long non-coding RNAs (lncRNAs) compared to WT controls. Predicted network analysis indicated that CD73KO specific lncRNAs control several signaling pathways. We found that genes in the JNK-MAPK-pathway were up-regulated upon infection in CD73KO and WT old but not in young mice. This corresponded to functional differences, as phosphorylation of the downstream AP-1 transcription factor component c-Jun was significantly higher in infected CD73KO and old mice neutrophils. Importantly, inhibiting JNK/AP-1 rescued the ability of these neutrophils to kill *S. pneumoniae*. Altogether, our findings revealed that neutrophils modify their gene expression to better adapt to bacterial infection and that this capacity declines with age and is regulated by CD73.

## Introduction

*Streptococcus pneumoniae* (pneumococcus) is an opportunistic pathogen that normally resides in the human nasopharynx but has the capacity to cause life-threatening infections that result in more than a million deaths annually (1). Pneumococcal infections are particularly a problem for elderly individuals. Despite the availability of vaccines and antibiotic therapies, *S. pneumoniae* remain a leading cause of community-acquired bacterial pneumonia in individuals above 65 years of age (2). According to a recent Active Bacterial Core surveillance report, individuals ≥50 years of age accounted for 71% of *S. pneumoniae* cases and 82% of associated deaths (3). Novel interventions are thus required to prevent a significant loss of life in the elderly and to combat the health and economic burden posed by this infection (4).

Neutrophils (also known as polymorphonuclear leukocytes or PMNs) play a central role in the clearance of *S. pneumoniae* infections. We and others found that PMNs are required for host resistance against pneumococcal infections (5–7) as depletion of PMNs prior to pneumococcal pulmonary challenge results in significantly higher bacteria burden in the lungs and increases lethality (7). It is well known that PMN antibacterial function declines with age (8, 9). We previously found that this could be recapitulated in mouse models where we observed a significant decrease in opsonophagocytic killing of *S. pneumoniae* by PMNs isolated from old mice compared to young controls (10). Strikingly, adoptive transfer of PMNs from young mice reversed the susceptibility of aged mice to pneumococcal pneumonia (10). This emphasizes the importance of PMNs in immunity and highlights their potential as targets for interventions that boost resistance of elderly hosts against infection. However, the host pathways that drive the age-associated decline in PMN function remain to be fully elucidated.

The extracellular adenosine (EAD) pathway plays an important role in host resistance to pneumococcal infection (7). Upon infection, ATP released by damaged or injured cells is converted into EAD by the sequential action of two extracellular enzymes: CD39 which converts ATP to AMP and CD73 which then dephosphorylates AMP to EAD (11). We previously found that genetic ablation or pharmacological inhibition of CD73 in mice results in higher pulmonary pneumococcal loads and systemic spread of infection (7). CD73 is required for the ability of PMNs to kill *S. pneumoniae* as PMNs isolated from young CD73KO mice fail to kill pneumococci *ex vivo* (7, 10, 12) . Importantly, age-driven changes in EAD pathway impair PMN anti-bacterial function. PMNs from old mice express significantly less CD73 than PMNs from young controls and supplementation with EAD reverses the age-driven decline in the ability of PMNs to kill *S. pneumoniae* (10).

The aim of this study was to investigate how aging impairs the antimicrobial activity of PMNs and what aspect of this is regulated by CD73. Although it was previously thought that PMNs are transcriptionally quiescent cells that kill bacteria with pre-packaged antimicrobial compounds, recent work has demonstrated that PMNs also undergo significant changes in their transcriptome in response to inflammation and bacterial infection (13, 14). Therefore, we examined global transcriptional changes in PMNs in response to *S. pneumoniae* infection *ex vivo* and how these responses are altered with aging and the absence of CD73. We found that infection with *S. pneumoniae* significantly altered the transcriptional profiles of PMNs from all host groups and that, importantly, active transcription was required for the ability of PMNs to kill bacteria. Surprisingly, we found that many more genes were down-regulated than up-regulated in response to infection. Down-regulated genes indicated a dampening of pro-inflammatory immune responses in PMNs from CD73KO and wild type (WT) old, but not in young hosts. Interestingly, higher numbers of long non-coding RNAs (lncRNAs) were found to be differentially expressed in PMNs from CD73KO mice compared to the PMNs from WT mice upon pneumococcal challenge. Predicted network analysis of these lncRNAs indicated that various immune signaling pathways are potentially regulated downstream of the EAD pathway. We also found an increased expression of Mitogen Activated Protein Kinase (MAPK) signaling pathway genes in PMNs from old and CD73KO but not young hosts. We confirmed that the activation of c-Jun N-terminal kinase/Activator protein-1 (JNK/AP-1), one of the MAPK-signaling pathways, was significantly up-regulated in PMNs from CD73KO and old mice compared to young controls in response to *S. pneumoniae* infection. Importantly, pharmacological inhibition of JNK/AP-1, reversed the defect in pneumococcal killing by PMNs from old and CD73KO mice, indicating that this pathway can potentially be targeted to reverse the age-related dysregulation of PMN responses.

## Materials and Methods

### Mice

Wild type (WT) young (4 months) and old (22-24 months) C57BL/6 mice were purchased from Jackson Laboratories (Bar Harbor, ME) and the National Institute on Aging colonies. CD73 knock-out (CD73KO) mice on a C57BL/6 background (15) were purchased from Jackson Laboratories and bred at a specific-pathogen free facility at the University at Buffalo. Young (4 months) CD73KO mice were used. Due to the limited availability of aged animals, male mice were used in all experiments. This work was performed in accordance with the recommendations from the Guide for Care and Use of Laboratory Animals published by the National Institutes of Health. All procedures were reviewed and approved by the Institutional Animal Care and Use Committee at the University at Buffalo.

### Bacteria

*S. pneumoniae* TIGR4 AC316 strain (serotype 4) (16) was a kind gift from Andrew Camilli. Bacteria were grown at 37°C in 5% CO_2_ in Todd-Hewitt broth supplemented with 0.5% yeast extract and oxyrase until cultures reached the mid-exponential phase. Aliquots were frozen at -80°C in growth media with 20% (v/v) glycerol. Aliquots were thawed on ice, washed, and diluted in PBS prior to use. Bacterial CFU were enumerated by serial dilution and dribble plating on TSA agar plates supplemented with 5% sheep blood (Northeast Laboratory).

### PMN isolation

Femurs and tibias of uninfected mice were flushed with RPMI 1640 supplemented with 10% FBS and 2 mM EDTA, and bone marrow cells were resuspended in PBS as described previously (12). PMNs were obtained through density gradient centrifugation using Histopaque 1119 and Histopaque 1077 as previously described (17). This method yields PMNs with 85-90% purity (12). To obtain ultrapure PMNs for RNA sequencing, negative selection EasySep Mouse Neutrophil Enrichment kit (StemCell#19762) was used following the manufacturer’s protocol. PMN purity was determined through flow cytometry and > 98% of cells were Ly6G+ (Fig S1).

### PMN infection and total RNA extraction

Ultrapure PMNs were isolated from young WT, old WT and young CD73KO mice. From each mouse, 10^6^ ultrapure PMNs were either infected with *S. pneumoniae* TIGR4 strain (MOI of 4) opsonized with 3% homologous mouse sera or mock-treated with 3% sera in buffer alone for 40 minutes at 37°C. Three mice per strain were used to obtain three distinct biological replicates of infected and mock-treated PMNs for a total of 18 samples. Following bacterial challenge, RNA was extracted from PMNs using RNeasy Mini Kit (Qiagen) as per the manufacturer’s protocol. TURBO deoxyribonucleic acid (DNA)-free kit (Invitrogen) was used to digest DNA from the samples. RNA concentrations and 260/280 ratios were determined using NanoDrop 1000 (Thermo Fischer Scientific).

### Illumina library preparation and RNA sequencing

Agilent 2100 Bioanalyzer was used to determine the integrity, purity and concentration of RNA samples. RNA integrity (RIN) score of 6.5 or above was considered acceptable for further analysis. Quality check revealed improper fragmentation of one sample (one mock-infected CD73KO sample), which was omitted from further analysis. Total RNA was enriched for mRNA using poly-(A)-selection (Illumina). NEB stranded RNA library prep kit (NEB) and NEB Ultra II RNA library prep kit (NEB) were used to prepare complementary DNA (cDNA) libraries for the remaining 17 samples, according to manufacturer’s protocol. RNA sequencing was carried out on an Illumina HiSeq2500 (Illumina) with a mid-output 75-cycle paired end with 10-20 million reads per sample at the Genomics and Bioinformatics core facility at the University at Buffalo. Details of the RNA samples along with RNA concentration and RIN score are provided in Supplementary Table I.

### Differential gene expression analysis

Per-cycle basecall (BCL) files generated by the Illumina HiSeq2500 were converted to per-read FASTQ files using bcl2fastq version 2.20.0.422 with default settings. FastQC version 0.11.5 was used to review the sequencing quality while FastQ Screen version 0.11.1 was used to determine any potential contamination. FastQC and FastQ Screen quality reports were summarized using MultiQC version 1.5 (18). Genomic alignments were performed using HISAT2 version 2.1.0 using default parameters (19). To differentiate between bacterial vs mammalian RNA, the resulting reads were aligned to NCBI GRCh38 as the reference genome. Sequence alignments were compressed and sorted into binary alignment map (BAM) files using samtools version 1.3. Counting of mapped reads for genomic features was performed using Subread FeatureCounts version 1.6.2 (20) (parameters:-s2–g gene_id –t exon –Q 60) and the annotation file specified with (–a) was the NCBI GRCh38 reference provided by Illuminas iGenomes. MultiQC software was used to summarize alignment as well as feature assignment statistics (18). Differentially expressed genes were detected using the Bioconductor package DESeq2 version 1.20.0 (21). Genes with one count or less were filtered out, and alpha was set to 0.05. Log2 fold-changes were calculated using DESeq2 using a negative binomial generalized linear models, dispersion estimates, and logarithmic fold changes integrated with Benjamini-Hochberg procedure to control the false discovery rate (FDR). A list of differentially expressed genes (DEGs) was generated through DESeq2. We defined a significant up- or down-regulation as a (fold change) ≥2 with FDR value < 0.05. The PCA plots were generated in ggplot2 package and the volcano plots were made using the Bioconductor package EnhancedVolcano.

### Gene ontology (GO) enrichment analysis

Functional enrichment analysis of significantly up- or down-regulated DEGs was performed on the Database for Annotation, Visualization and Integrated Discovery (DAVID) (22) using the default settings. For each comparison, gene functions were categorized into biological process, molecular function, and cellular components. These gene functions were analyzed separately for up- or down-regulated DEGs. DAVID was used to further perform pathway analysis and to retrieve pathway maps based on the identified DEGs. All functional categories and pathways with *p*-value < 0.05 were considered significant. The complete data are available as Supplementary material and on NCBI website with accession number GSE150811.

### LncRNA network analysis

In order to elucidate the possible function and biological process of long non-coding RNAs (lncRNAs) identified in this screen, we performed computational prediction of the potential lncRNA-target interaction. LncRNAs bind to complementary sequence of neighboring or target genes to repress expression. Thus, if a lncRNA is up-regulated, it is predicted to down-regulate the expression of the target gene and vice versa. We performed computational prediction of lncRNA-target interactions using LncTar software for prediction of lncRNA-RNA interactions through free energy minimization. Using the normalized binding free energy (ndG), we selected a value of -0.02 as cutoff for the analysis as previously described (23). In order to confirm the reliability of our prediction analysis, we further used LncRRIsearch, an online server for prediction of lncRNA-target interaction to validate the result from the previous analysis. Briefly, we searched the genomic location of all our lncRNAs from the mouse genome (GRCm38.p6) and nucleotide sequences of the lncRNAs and their neighboring genes were retrieved for prediction of potential lncRNA-RNA interactions. In order to gain understanding of the possible biological process and physiological pathways, we catalogued all potential target genes and performed functional enrichment analysis to identify significantly affected pathways using a combination of gene ontology (GO) term, PANTHER and KOBAS (http://kobas.cbi.pku.edu.cn/kobas3) as previously described (24, 25). Networks were then generated indicating the likelihood of the focus lncRNAs, gene targets and biological process in the network being found together by chance including concomitant lncRNAs co-regulating one or more targets (26). The networks, pathways, and biological functional classification were generated using Cytoscape version 3.7.2.

### RNA sequencing data accession number

The data presented and discussed in this manuscript along with all the RNA sequencing files and raw data files have been deposited in the NCBI’s Gene Expression Omnibus (GEO), and is accessible through GEO Series accession number GSE150811 (https://www.ncbi.nlm.nih.gov/geo/query/acc.cgi?acc=GSE150811).

### Real time PCR

RT-PCR was used to validate the expression of some of the differentially expressed genes identified in RNA sequencing. For this, the same RNA samples previously used to prepare Illumina libraries were used. Following treatment with indicated inhibitors and challenge with *S. pneumoniae*, RNA was extracted from 1 x 10^6^ PMNs/condition and DNA digested as described above. For all RT-PCR reactions, 500 ng of each sample was converted into cDNA using Super-Script VILO^TM^ cDNA synthesis kit (Life Technologies) according to the manufacturer’s protocol. RT-PCR was performed using CFX96 Touch ^TM^ Real-Time PCR Detection System from Bio-Rad. CT (cycle threshold-values) were determined using the following TaqMan probes from Life Technologies (Thermo Fischer Scientific): GAPDH (Mm99999915_m1), IL-10 (Mm01288386_m1), c-FOS (Mm00487425_m1), Cybr4 (Mm01144487_m1), Hsp72 (Mm01159846_s1), Rgl1 (Mm00444088_m1), ADOR2B (Mm00839292_m1), Rrad (Mm00451053_m1), Tnip1 (Mm01288484_m1), DUSP1 (Mm00457274_g1), c-JUN (Mm00495062_s1), Nr4a1 (Mm01300401_m1), Sifn1 (Mm00624380_m1), Tubb6 (Mm00660543_m1), and Atf3 (Mm00476032_m1). All samples were run in duplicates. Data were analyzed by the comparative threshold cycle (2^-ΔCT^) method, normalizing the CT values obtained for target gene expression to those for GAPDH of the same sample. For comparison of expression levels upon infection, relative quality of transcripts (RQ) were calculated by the ΔΔCT method by using the formula RQ = 2-^(ΔΔCT)^ (27). ΔΔCT values were obtained by using the formula ΔΔCT = ΔCT_infected_ – ΔCT_uninfected_.

### Opsonophagocytic killing assay (OPH)

The ability of PMNs to kill *S. pneumoniae* ex vivo was measured using a well-established OPH killing assay as previously described (7, 10, 12, 28). Briefly, 1×10^5^ PMNs were incubated with 1×10^3^ bacteria grown to mid-log phase and pre-opsonized with 3% mouse sera in 100μl reactions of HBSS containing 0.1% gelatin. Reactions were then rotated at 37°C for 45 minutes. Where indicated, PMNs were incubated with Actinomycin D (transcription inhibitor), Cycloheximide (translation inhibitor), Anisomycin (JNK stimulator), SR11302 (AP-1 inhibitor), JNK-IN-8 (JNK inhibitor), or HBSS (vehicle control) for 30 minutes prior to infection. Anisomycin and SR11302 were purchased from Tocris Biosciences and Actinomycin D, Cycloheximide and JNK-IN-8 from Sigma. Percent killing was determined by dribble plating on blood agar plates and calculated in comparison to the no PMN control under the same conditions (+/− treatments).

### Phosphorylated c-Jun measurement

The ability of *S. pneumoniae* to induce phosphorylation of c-Jun was measured by flow cytometry. Briefly, 5×10^5^ PMNs were challenged with pre-opsonized *S. pneumoniae* TIGR4 at MOI of 4 in 100μl reactions of HBSS containing 0.1% gelatin. Reactions were then rotated at 37°C for indicated time points. Where indicated, PMNs were incubated with, Anisomycin (JNK stimulator), SR11302 (AP-1 inhibitor), JNK-IN-8 (JNK inhibitor), or HBSS (vehicle control) for 30 minutes prior to infection. Following incubation, cells were fixed with Cytofix (BD Bioscience) and permeabilized by ice cold methanol. Cells were then stained for fluorophore-tagged antibodies against Ly6G (BD Bioscience # 5605991), phospho c-Jun (Ser73) (Cell Signaling # 12714S) (29) and total c-Jun (Cell Signaling # 15683S) at 1:50 dilutions per manufacturer’s protocol. Fluorescence intensities were measured on a BD Fortessa and at least 10,000 events were analyzed using FlowJo.

### Flow cytometry

Anti-Ly6G (IA8, BioLegend) antibodies were used to determine the purity of isolated PMNs. Staining was performed in the presence of Fc-block (BD Bioscience). Fluorescence intensities were measured on a BD Fortessa and at least 2,000 events were analyzed using FlowJo.

### Statistics

OPH and flow cytometry data were analyzed using Prism8 (Graph Pad). Bar graphs represent the mean values +/- SD. 1-sample t-test or Student’s t-test were used to determine significant differences as indicated. Correlation of mRNA expression by RNA-Seq and qPCR was assessed by Pearson correlation analysis. All *p*-values less than 0.05 were considered significant (as indicated by asterisks).

## Results

### Active transcription and translation are important for the ability of PMNs to kill *S. pneumoniae*

We previously reported that PMNs from old mice fail to efficiently kill *S. pneumoniae*, in part due to a decline in the surface expression of CD73 and extracellular adenosine production (10). In this study, we wanted to explore whether CD73 and age-driven changes in the transcriptome impair PMN antimicrobial function. As PMNs are known to have antimicrobial products pre-synthesized and packaged during maturation for rapid immune response (30), we investigated the importance of active transcription and translation in the ability of PMNs to kill *S. pneumoniae.* To do this, we used a well-established *ex vivo* opsonophagocytic killing assay (7, 31) where we isolated PMNs (Ly6G^+^) from the bone marrow of young C57BL/6 wild type (WT) mice and treated them with either Actinomycin D (transcription inhibitor (32)) or Cycloheximide (translation inhibitor (33)) at concentrations that do not impair cellular viability (32, 34) prior to infection with *S. pneumoniae*. We found that treating PMNs with Actinomycin D caused a significant 2-fold decrease in bacterial killing compared to vehicle control (VC), while treatment with Cycloheximide completely abrogated the ability of PMNs to kill bacteria and instead enabled bacterial growth in the presence of PMNs (Fig. 1A). These findings suggest that active transcription of new mRNAs and formation of new proteins is crucial for optimal anti-pneumococcal responses.

**Figure 1.**
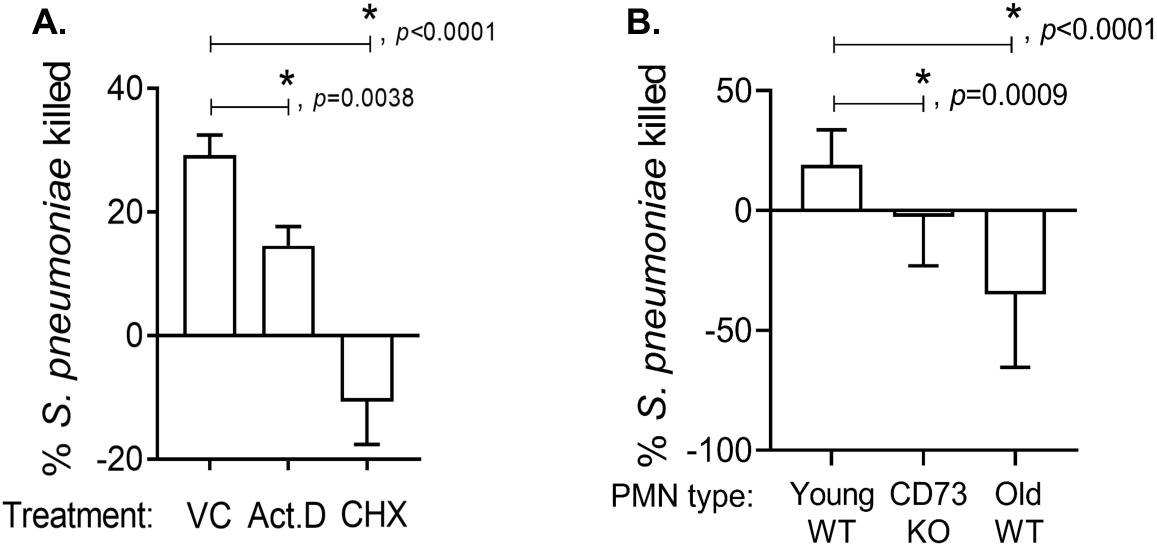
Active transcription and translation are required for the ability of PMNs to kill *S. pneumoniae.* (A) PMNs isolated from the bone marrow of C57BL/6 young WT mice were treated with 5µg/mL of Actinomycin D (Act.D) or 10µg/mL of Cycloheximide (CHX), or PBS (vehicle control) for 30 minutes at 37°C. Treated neutrophils were then infected with *S. pneumoniae* TIGR4 pre-opsonized with homologous sera for 45 minutes at 37°C. Reactions were plated on blood agar plates and the percentage of bacteria killed compared to a no PMN control under the same condition was calculated. Positive percent killing indicates bacterial death while negative percent indicates bacterial growth. (B) PMNs isolated from the bone marrow of C57BL/6 young WT, old WT and CD73KO mice were infected with *S. pneumoniae* TIGR4 pre-opsonized with homologous sera for 45 minutes at 37°C. Reactions were plated on blood agar plates and the percentage of bacteria killed compared to a no PMN control under the same condition was calculated for each strain. (A and B) Data shown are pooled from six separate experiments (n=6 biological replicates) where each condition was tested in triplicate (n=3 technical replicates) per experiment. Asterisks indicate significant differences between the indicated groups as calculated by Student’s t-test.

### Profiling of mRNA expression

We wanted to test whether there are age-related changes in mRNA expression that renders PMNs ineffective in their antimicrobial function. In addition, we were interested in investigating whether any of the age-driven changes were shared by PMNs that lack CD73. We first re-confirmed that aging and lack of CD73 significantly blunts the ability of PMNs to kill *S. pneumoniae ex vivo* (Fig. 1B). Next, RNA sequencing was used to compare the transcriptional profiles of PMNs from young WT, old WT, and young CD73KO mice at baseline and upon infection. For RNA isolation, we obtained an ultrapure PMN population (approximately 99% purity, Fig S1) from the bone marrow of mice using negative selection (see materials and methods). Three mice were used per strain. Efficient killing of pneumococci by PMNs from young controls requires opsonization (35). Therefore, to more closely mimic *in vivo* conditions and the opsonophagocytic killing assay (Fig. 1B), PMNs isolated from each mouse were either challenged with *S. pneumoniae* TIGR4 strain (at a multiplicity of infection or MOI 4) opsonized with homologous mouse sera from the same mouse for 40 minutes or mock-challenged with sera containing buffer. We focused on the 40-minute time point as this is a standard time used in *ex vivo* killing assays (7, 10, 12) and it allows us to examine differences in antimicrobial function (Fig. 1B), while maintaining PMN viability (≤ 20% PMN necrosis (PI+), Fig. S4B). Detailed methods on ultrapure PMN isolation and subsequent RNA sequencing workflow are in the materials & methods section and summarized in Fig. 2A. Differentially expressed genes (DEGs) were analyzed using DESeq2 and significant differential expression of a gene was defined as expression with fold change value of ≥ 2.0 and a false discovery rate (FDR) < 0.05.

**Figure 2.**
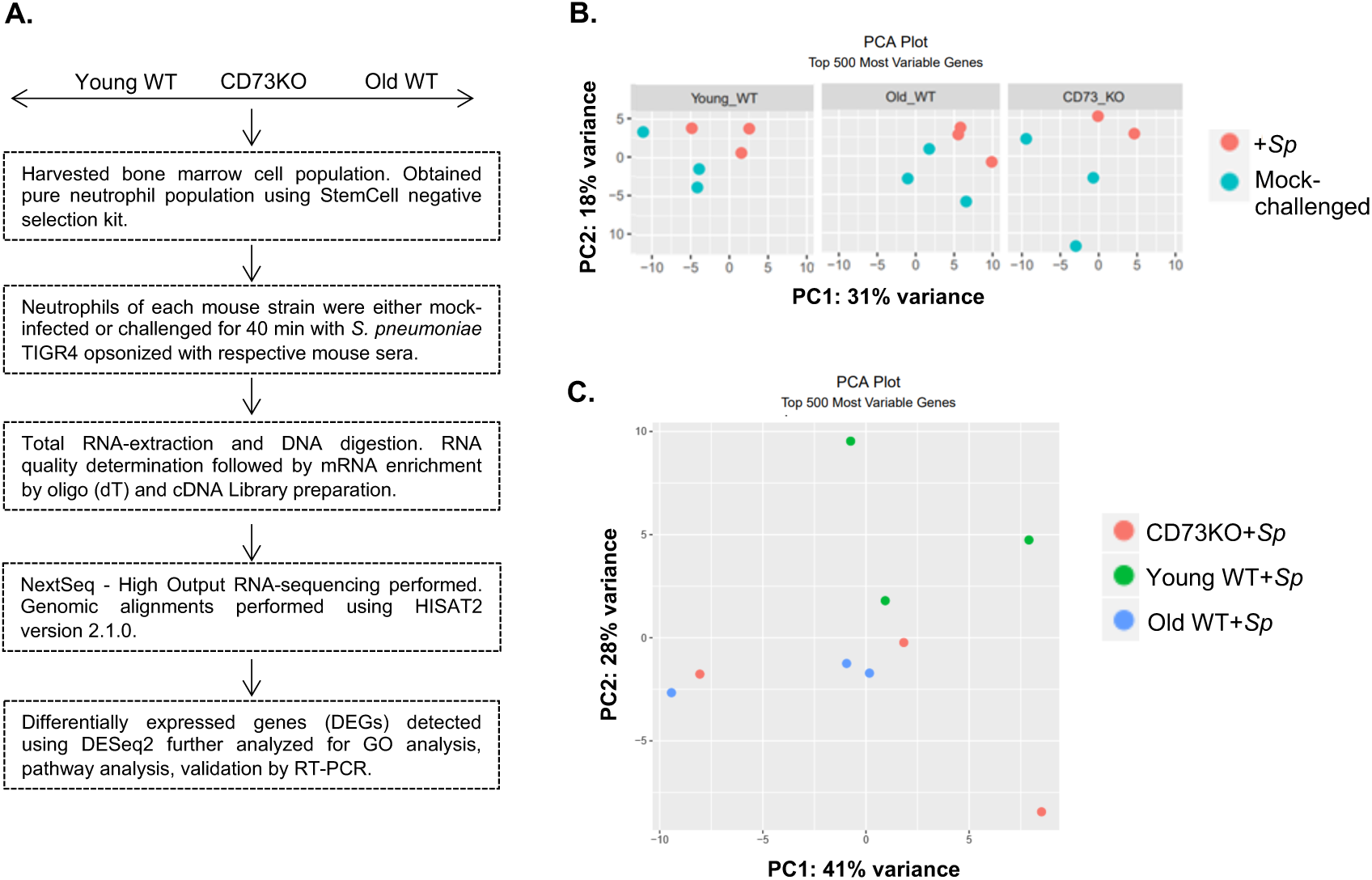
RNA sequencing experimental approach. (A) Schematic diagram of PMN isolation, sample preparation, and RNA sequencing analysis. (B and C) Principal component analysis (PCA) plot showing variance in mRNA expression (post data normalization) in un-infected or *S. pneumoniae* challenged samples, presented as separate plots for each mouse strain (B) or all infected samples on the same plot (C).

### Mock-infected PMNs from young, old and CD73KO mice show limited differences in mRNA profiles

To determine if there is an intrinsic age-driven change in expression of genes that shape antimicrobial responses, we compared mRNA expression profiles of mock-challenged PMNs from old WT mice to that of young WT controls. Keeping the expression of PMNs from young WT mice as baseline, we found a total of 23 DEGs to be up-regulated in PMNs from old mice (Table I). Surprisingly, 15 of these DEGs corresponded to the category of either Immunoglobulin heavy chain variable regions or Immunoglobulin kappa chain variable region (Table I). mRNA levels of certain variable region genes have been previously shown to vary in PMNs, although these cells do not express immunoglobulins (36). PMNs from old WT mice also showed up-regulation of a few other genes including *Calca* (calcium regulation and cAMP activity), *Mt2* (metal ion regulation), *Ces1d* (lipase activity), *Col5a1* (type V collagen) and *C130026l21Rik* (unannotated lncRNA) (Table I). Interestingly, none of the genes known for their role in PMN antimicrobial function showed an age-driven differential expression under baseline conditions.

**Table I.**
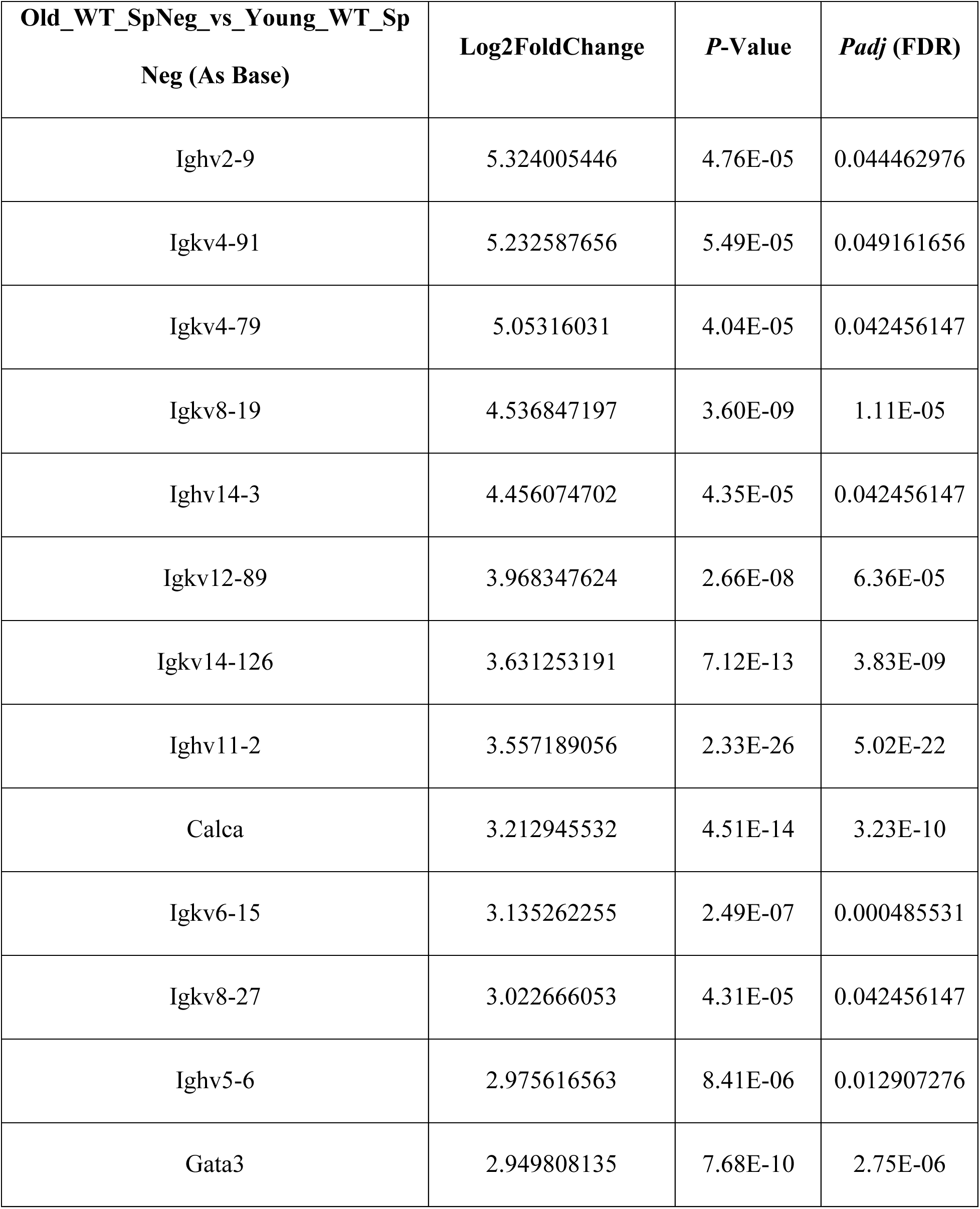
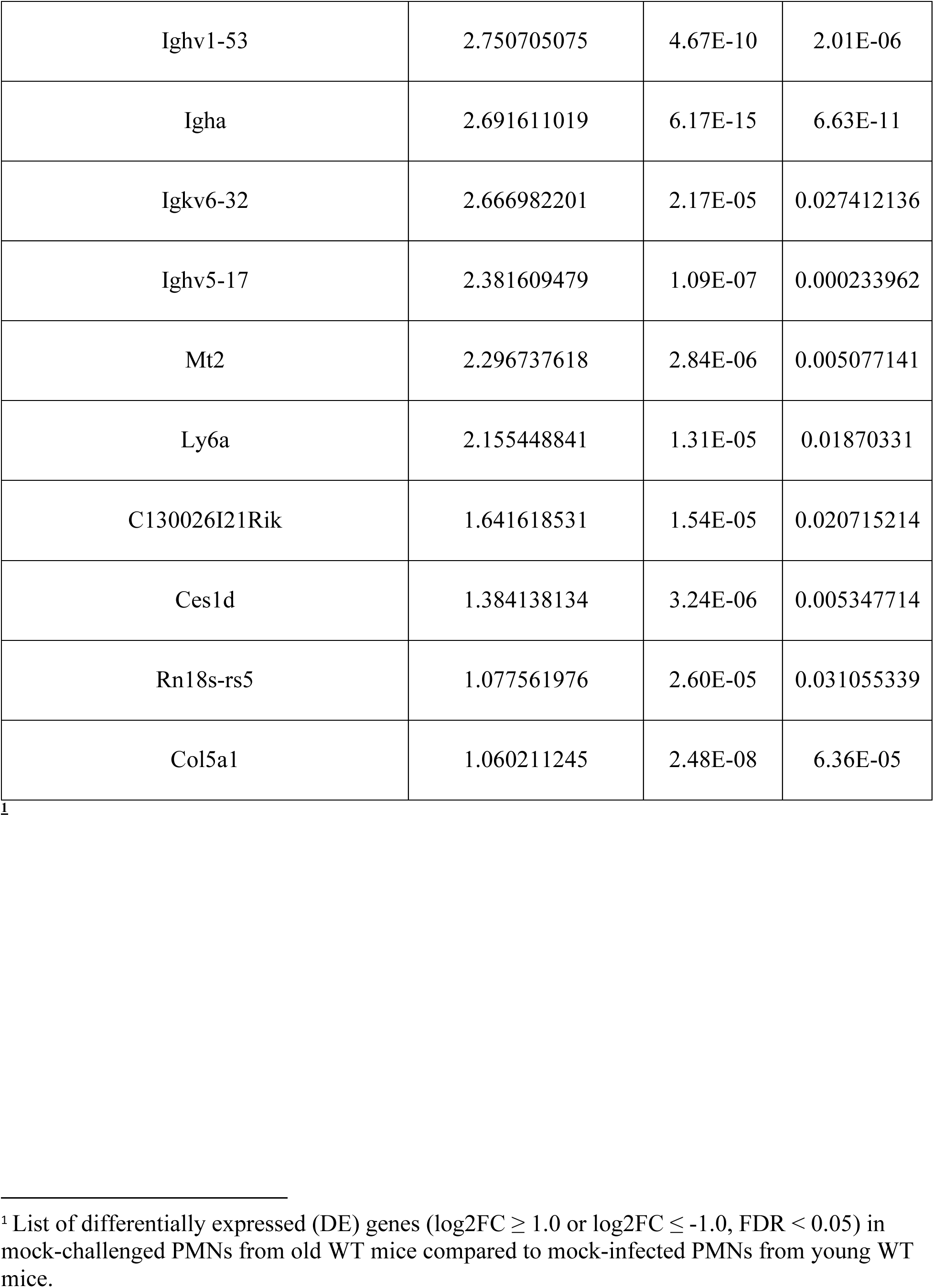
Differentially expressed genes in mock-challenged PMNs from old WT mice compared to young WT mice.

To determine if there is an intrinsic CD73-driven change in expression of genes that shape antimicrobial responses, we then compared mRNA expression profiles of mock-stimulated PMNs from young CD73KO mice to that of young WT mice. As shown in Table II, we noted that only 8 genes that were differentially expressed in resting PMNs with an equal number of up-regulated and down-regulated DEGs. Up-regulated DEGs included *Gm11868* (a lncRNA with predicted histone demethylase activity in *Drosophila*), *Gm13456* (a pseudogene related with somatic muscle development activity in *Drosophila*), *Gm6548* (unannotated pseudogene), and *Ighv9-4* (corresponds to the category of Immunoglobulin heavy chain variable region). As expected, the down-regulated DEGs included *NT5E* (that encodes for CD73). Other down-regulated DEGs included *Fam63b* (ubiquitin carboxyl-terminal hydrolase activity), *Aqp9* (transmembrane transporter activity) and *Cyb5r4* (NADPH-cytochrome reductase activity). As observed in old mice, none of the known antimicrobial genes showed a differential expression in mock-challenged CD73KO PMNs. In summary, we found limited differences in mRNA expression in mock-challenged PMNs from WT and CD73KO mice as well as across host age.

**Table II.** Differentially expressed genes in mock-challenged PMNs from CD73KO mice compared to young WT mice.

| **CD73KO_SpNeg_vs_Young_WT_Sp**<br>**Neg (As Base)** | **Log2FoldChange** | ***P*-Value** | ***P*adj (FDR)** |
| --- | --- | --- | --- |
| Gm11868 | 5.382471685 | 4.98E-30 | 3.69E-26 |
| Ighv9-4 | 5.305558233 | 1.53E-06 | 2.84E-03 |
| Gm13456 | 2.127675061 | 1.06E-32 | 1.57E-28 |
| Gm6548 | 1.275332385 | 2.14E-07 | 4.52E-04 |
| Fam63b | -1.142265331 | 5.00E-21 | 2.47E-17 |
| Aqp9 | -1.14542389 | 8.13E-06 | 1.20E-02 |
| Cyb5r4 | -1.393240659 | 9.15E-17 | 2.71E-13 |
| Nt5e | -2.880728481 | 2.53E-18 | 9.39E-15 |

<sup>1</sup> List of differentially expressed (DE) genes ( $\log_2FC \geq 1.0$ or $\log_2FC \leq -1.0$ , $FDR < 0.05$ ) in mock-challenged CD73KO PMNs compared to mock-infected WT PMNs from young mice.

### *S. pneumoniae* induces global changes in transcriptome profiles

We next wanted to determine whether *S. pneumoniae* infection induced any transcriptional changes in PMNs. To do that, the global transcriptome profiles of infected and mock-challenged PMNs were characterized for each mouse group. Principal Component Analysis (PCA) was done prior to and after pneumococcal infection to investigate changes in patterns of mRNA expression between the different groups. We found that infection with *S. pneumoniae* resulted in major transcriptome changes in all three PMN types (Fig. 2B). Analysis of PMNs from each mouse group clearly showed distinct patterns of mRNA expression between the mock-infected and infected samples with combined total variance of 49% (PC1 and PC2), suggesting a distinct response of PMNs to *S. pneumoniae*. In PMNs from all three mouse groups, the mock-challenged samples formed a cluster separate from the corresponding infected samples (Fig. 2B). When we compared infected PMNs across the different mouse groups, we found that while CD73KO PMNs showed variation, PMNs from young WT mice clustered distinctly from the corresponding old mice (Fig. 2C). In summary, infection with *S. pneumoniae* triggers global changes in PMN transcriptome profiles that differed across host age.

### Genes and functional categories up-regulated in response to *S. pneumoniae*

We then explored the genes whose expression was up-regulated upon PMN infection and how this varied among the different host groups. By selecting DEGs with at least 2-fold change in expression compared to mock-infected controls for each mouse group, we surprisingly found only a small number of genes (10 per group) that were up-regulated in PMNs from WT mice in response to pneumococcal challenge, regardless of age (Fig. 3A and 4A). In contrast, CD73KO PMNs showed the strongest transcription response to bacterial infection with 36 up-regulated genes (Fig. 5A). Some of the up-regulated DEGs were common in PMNs from all three mouse groups (Fig. 6A). The six overlapping DEGs (*Osm, Fos, Jun, Zfp36, Egr1 and Atf3*) belonged to the categories of growth regulators, transcription factors and transcription and translation regulators. When we compared DEGs that were commonly up-regulated in PMNs from old WT and CD73KO, but not in young WT mice, we found only two DEGs (Fig. 6A): *Slfn1* that has a known role in cell proliferation and immune response and *Nr4a1*, a transcription factor. When examining the DEGs that were up-regulated in response to infection that were specific to CD73KO PMNs, we found increased expression of *Btg2* (regulation of cell cycle), *Zcchc4* (nucleic acid binding and methyltransferase activity), *Dusp1* (phosphatase activity), *Klhl42* (ubiquitin-protein transferase activity), *Snai1* and *Hlx* (sequence specific DNA binding activity), *F3* (phospholipid binding and cytokine receptor activity), *Hspa1a* and *Hspa1b* (ubiquitin protein ligase binding and protein folding chaperone), *Tacstd2* (calcium signaling), and *Rhob* (GDP and GTP binding activity). A number of up-regulated DEGs from CD73KO PMNs belonged to the category of lncRNAs that have not been functionally annotated, thus their roles in cellular function are currently unknown.

**Figure 3.**
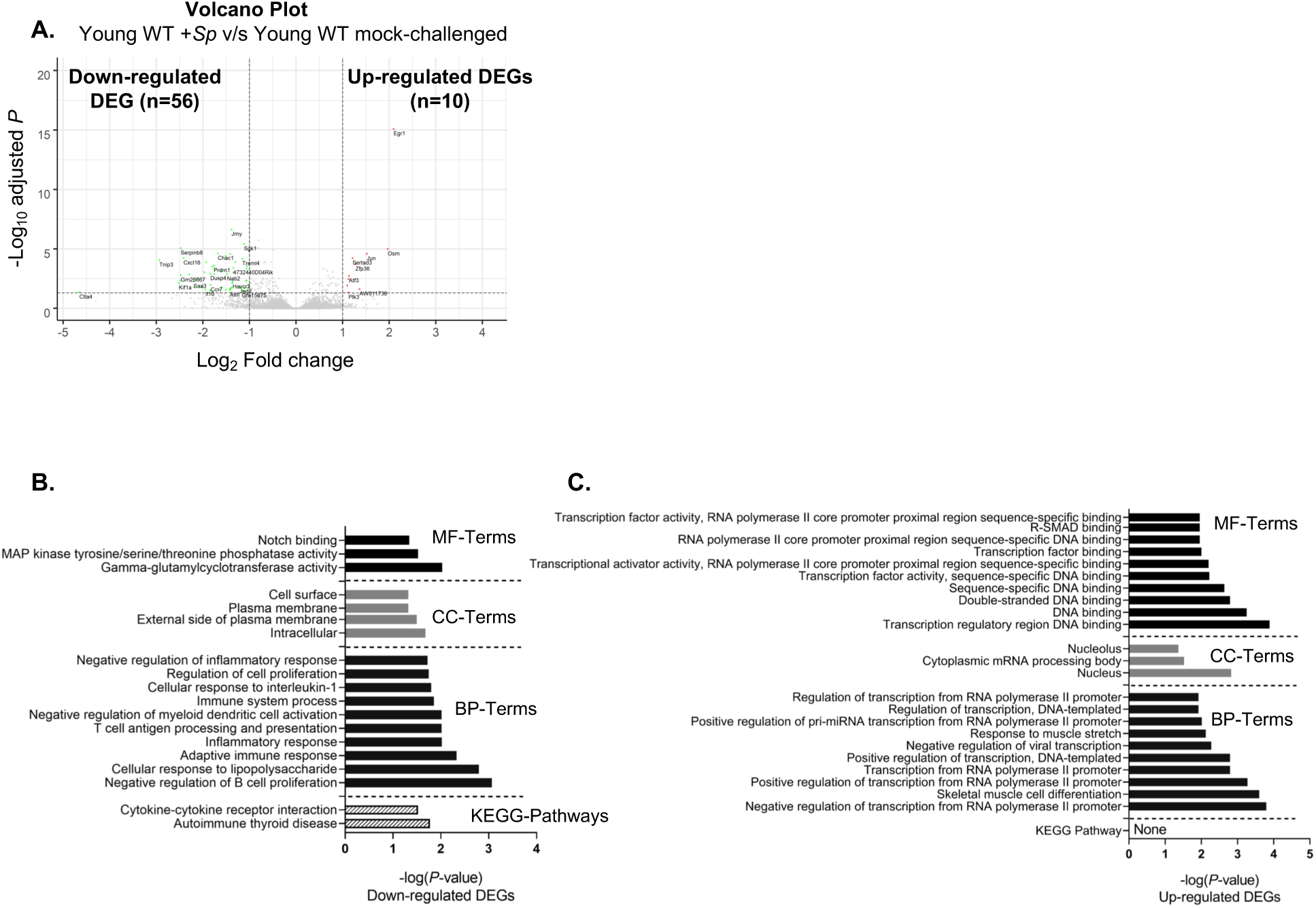
Analysis of differentially expressed genes in PMNs from young WT mice in response to *S. pneumoniae* infection. (A) Volcano plot representing differential gene expression (DEG) (FDR <0.05) in PMNs isolated from the bone marrow of young WT mice in response to *ex vivo* challenge with *S. pneumoniae* TIGR4 compared to mock-challenged control. Genes marked in green represent significantly down-regulated DEGs (log2FC ≤ -1.0, FDR < 0.05) and genes marked in red represent significantly up-regulated DEGs (log2FC ≥ 1.0, FDR < 0.05). (B and C) Gene Ontology (GO) enrichment analysis using DAVID indicating the top 10 significant (*p* ≤ 0.05) Biological Process (BP), Molecular Function (MF), Cellular Component (CC) and KEGG Pathway terms for significantly down-regulated DEGs (B) and significantly up-regulated DEGs (C).

**Figure 4.**
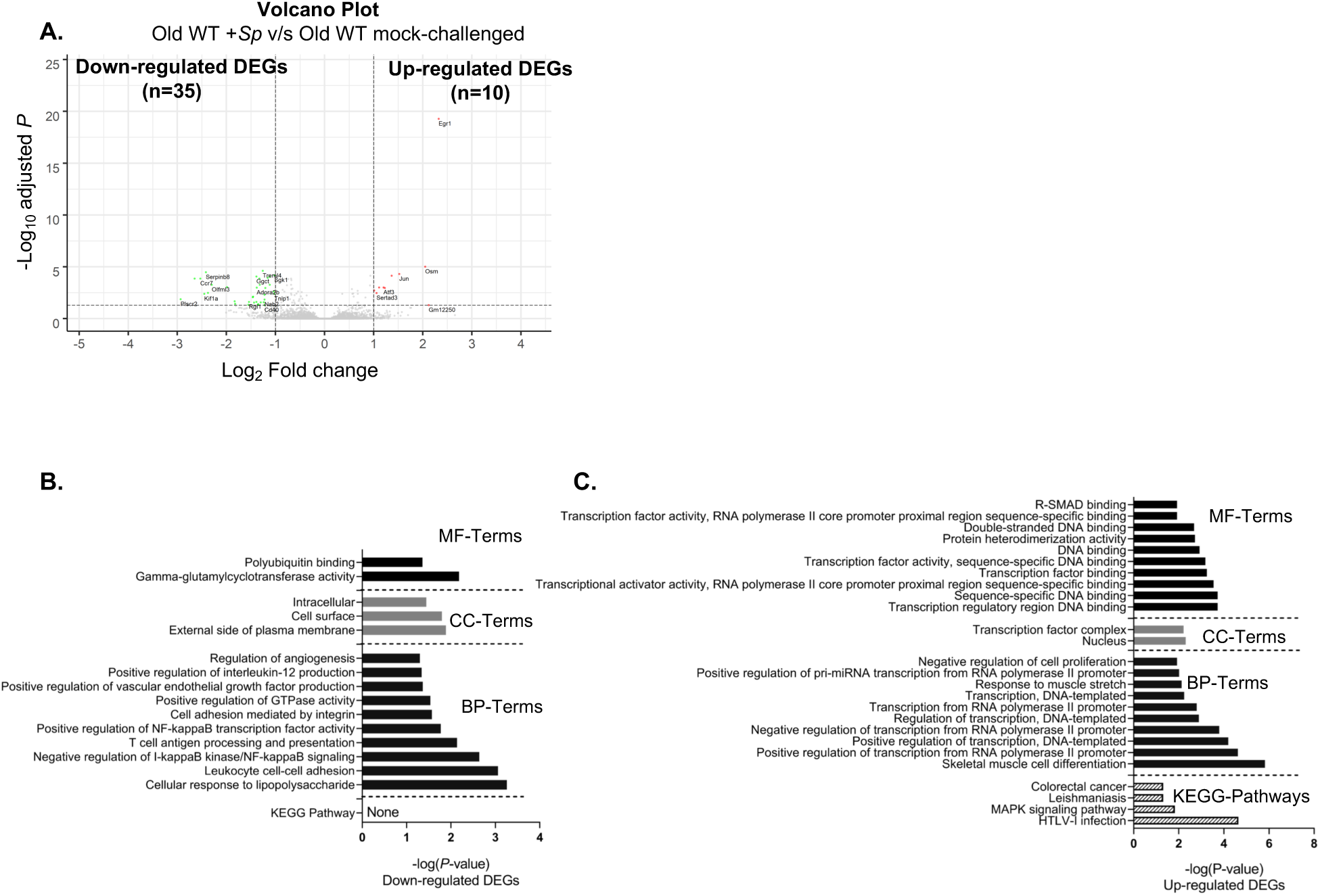
Analysis of differentially expressed genes in PMNs from old WT mice in response to *S. pneumoniae* infection. (A) Volcano plot representing differential gene expression (DEG) (FDR <0.05) in PMNs isolated from the bone marrow of old WT mice in response to *ex vivo* infection with *S. pneumoniae* TIGR4 compared to mock-infected control. Genes marked in green represent significantly down-regulated DEGs (log2FC ≤ -1.0, FDR < 0.05) and genes marked in red represent significantly up-regulated DEGs (log2FC ≥ 1.0, FDR < 0.05). (B and C) Gene Ontology (GO) enrichment analysis using DAVID indicating the top 10 significant (*p* ≤ 0.05) Biological Process (BP), Molecular Function (MF), Cellular Component (CC) and KEGG Pathway terms for significantly down-regulated DEGs (B) and significantly up-regulated DEGs (C).

**Figure 5.**
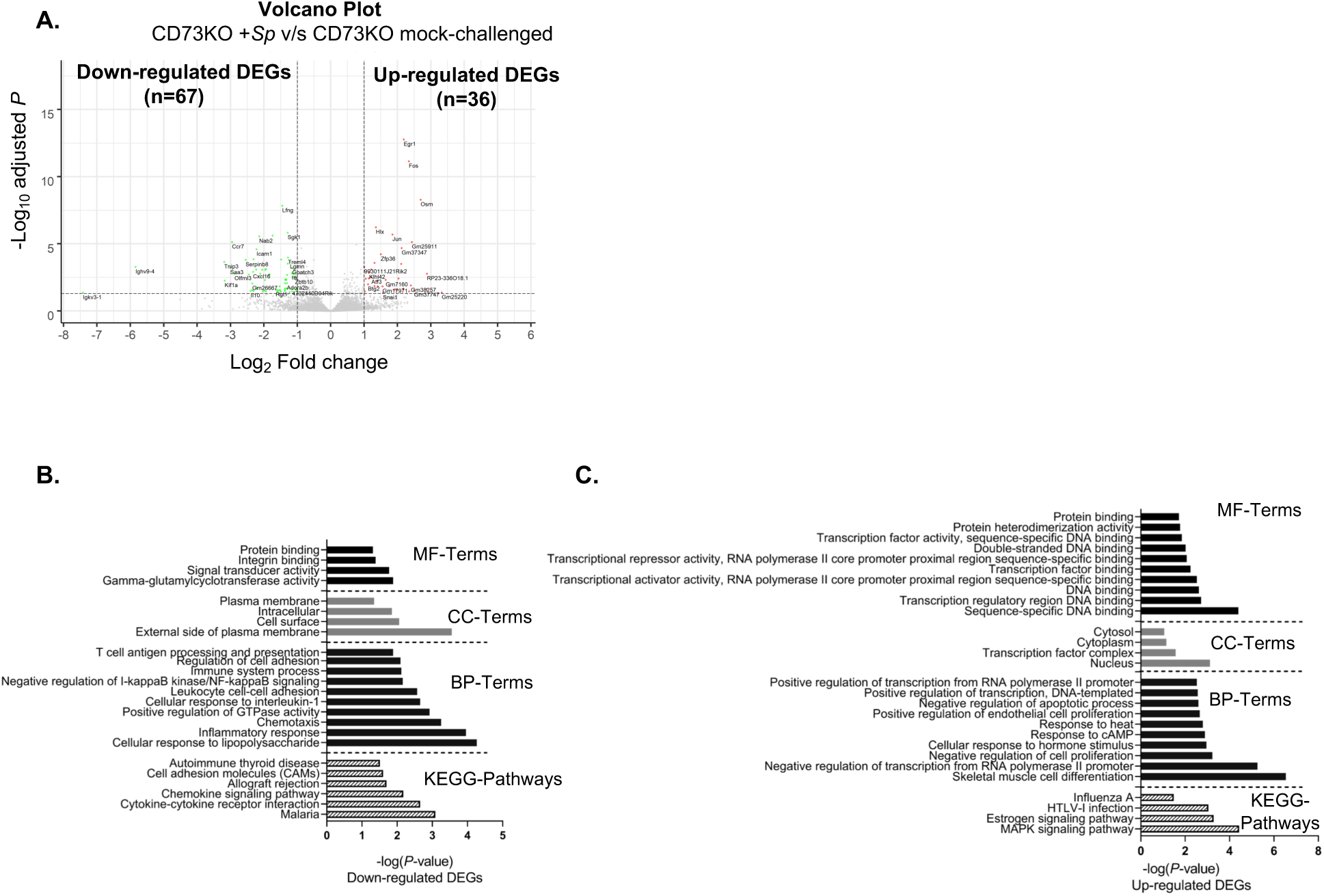
Analysis of differentially expressed genes in PMNs from young CD73KO mice in response to *S. pneumoniae* infection. (A) Volcano plot representing differential gene expression (DEG) (FDR <0.05) in PMNs isolated from the bone marrow of CD73KO mice in response to *ex vivo* challenge with *S. pneumoniae* TIGR4 compared to mock-infected control. Genes marked in green represent significantly down-regulated DEGs (log2FC ≤ -1.0, FDR < 0.05) and genes marked in red represent significantly up-regulated DEGs (log2FC ≥ 1.0, FDR < 0.05). (B and C) Gene Ontology (GO) enrichment analysis using DAVID indicating the top 10 significant (*p* ≤ 0.05) Biological Process (BP), Molecular Function (MF), Cellular Component (CC) and KEGG Pathway terms for significantly down-regulated DEGs (B) and significantly up-regulated DEGs (C).

**Figure 6.**
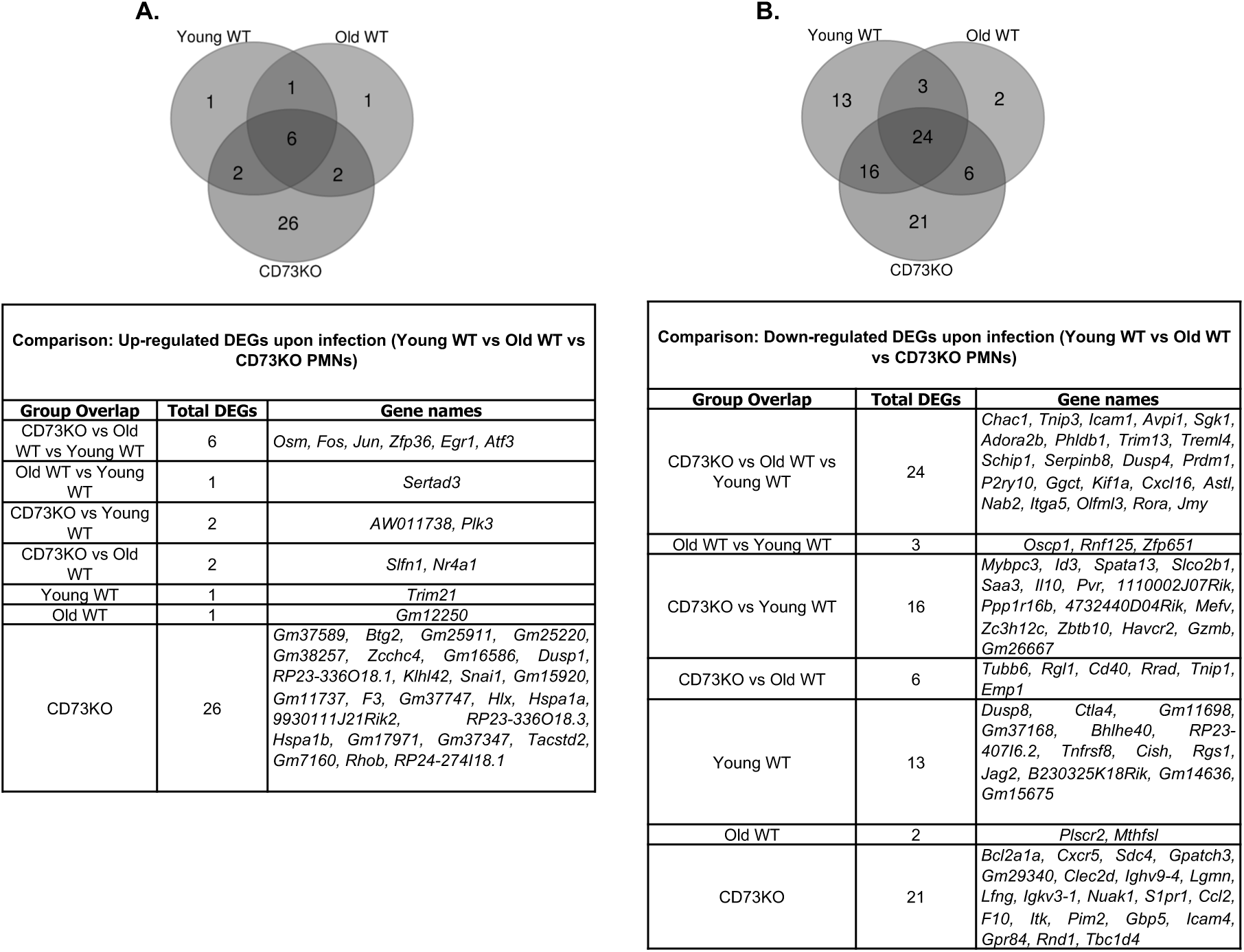
Venn diagrams showing distribution of significantly up-regulated or down-regulated genes across host groups in response to *S. pneumoniae* infection. Distribution of significantly up-regulated (log2FC ≥ 1.0, FDR < 0.05) (A) and significantly down-regulated (log2FC ≤ -1.0, FDR < 0.05) (B) DEGs in PMNs from young WT vs old WT vs CD73KO mice upon pneumococcal challenge.

We next grouped up-regulated genes into different functional categories (Supplementary Tables II, IV and VI). Overall, there was a significant overlap in the annotated processes between PMN from the three mouse groups with DEGs falling mainly into the categories of DNA binding, transcription regulation and transcription factor activity (Fig. 3C, 4C and 5C) as many of these DEGs are known to regulate gene expression either as co-activators, regulators or transcription factors (*Fos, Jun, Egr1, Atf3, Sertad3, Nr4a1, F3* and *Hlx*). These data indicate that upon challenge with *S. pneumoniae*, PMNs may have undergone transcriptional reprogramming as indicated by up-regulation of genes involved in transcription activation or transcription regulation.

### Genes and functional categories down-regulated in response to *S. pneumoniae*

Genes whose expression was down-regulated upon PMN infection were examined including their variation among the different host groups. Interestingly, we found more genes (2-3-fold more) that were down-regulated than up-regulated in PMNs in response to infection in all mouse groups (Fig 3A, 4A, and 5A). A total of 56 genes were down-regulated in PMNs from young mice, while only 35 genes were down-regulated in PMNs from old mice in response to pneumococcal challenge (Fig. 3A and 4A). As observed with the up-regulated DEGs, CD73KO PMNs showed strongest transcriptional response following *S. pneumoniae* challenge with 67 down-regulated DEGs (Fig. 5A). Overall, there was considerable overlap observed between PMN from the three mouse groups (Fig. 6B). The 24 overlapping DEGs belong to categories of immune and inflammatory response (*Tnip3, Icam1, Sgk1, Prdm1, Cxcl16*, and *Prdm1*), MAPK-signaling (*Dusp4*), cell-surface signaling (*Adora2b, Treml4, P2ry10,* and *Itga5*), transcription regulation (*Jmy, Rora* and *Nab2*), microtubule organization (*Kif1a*), protein regulation (*Trim13*), cell cycle and cell-cell adhesion (*Avp1* and *Serpinb8*), actin cytoskeleton (*Phldb1*), podocyte function (*Schip1*), apoptosis (*Ggct*), metallopeptidase (*Astl*), embryonic development function and tumorigenesis (*Olfml3*) and Notch-signaling (*Chac1*). Comparison of DEGs that were commonly down-regulated in PMNs from old WT and CD73KO mice showed 6 overlapping down-regulated DEGs that were not differentially expressed in PMNs from young WT mice (Fig. 6B). These included *Tubb6* (microtubule organization)*, Rgl1* (guanine nucleotide exchange factor)*, Rrad* (GTPase activity)*, Cd40* (immune and inflammatory response)*, Tnip1* (inflammatory response) and *Emp1* (cell-cell interaction and cell proliferation). These findings point towards an overall age-related decrease in immune and inflammatory response, characteristics of which are also shared by CD73KO PMNs.

To further understand how CD73 regulates the transcriptional profile during infection, we examined the distribution of DEGs that were only down-regulated in CD73KO, but not in WT PMNs (Fig. 6B). These included migration related genes *Cxcr5* (C-X-C-chemokine receptor activity), *Cccl2* (CCR2 chemokine receptor binding), and *Icam4* (integrin binding); G-protein coupled receptors related genes *S1pr1* (G protein-coupled receptor binding) and *Gpr84* (G protein-coupled peptide receptor activity); GTP related genes *Gbp5* (GTP hydrolysis), *Rnd1* (GTPase activity), and *Tbc1d4* (GTPase activator activity); kinase related genes *Sdc4* (protein kinase C binding), *Itk* (Tyrosine kinase activity), *Nuak1* and *Pim2* (serine/threonine protein kinase activity); and genes involved in other processes *Bcl2a1a* (apoptotic process), *Gpatch3* (nucleic acid binding), *Clec2d* (transmembrane signaling receptor activity), *Lgmn* (endopeptidase activity), *Lfng* (acetylglucosaminyl transferase activity) and *F10* (calcium and phospholipid binding). These data suggest potential dysregulation in PMN migration in response to *S. pneumoniae* in the absence of CD73, which is consistent with our previous findings (7).

To elucidate the PMN responses dampened upon pneumococcal challenge in susceptible vs. resistant hosts, we compared the distribution of DEGs that were down-regulated in PMNs from young WT mice only but not in PMNs from old WT or CD73KO PMNs. These included *Dusp8* (tyrosine/serine/threonine phosphatase activity), *Ctla4* (negative regulator of T-cell responses), *Bhlhe40* (transcriptional repressor activity), *Tnfrsf8* (transmembrane signaling receptor activity), *Cish* (1-phosphatidylinositol-3-kinase regulator activity), *Rgs1* (GTPase activator activity) and *Jag2* (calcium ion binding and growth factor activity). These data suggest that select genes that inhibit immune responses are down-regulated in young WT hosts to better respond to *S. pneumoniae* challenge.

We further categorized down-regulated genes into different functional categories (Supplementary Tables III, V and VII). As expected, many genes were shared by more than one functional category. Overall, the functional categories which were commonly down-regulated in all three PMN types included cellular response to lipopolysaccharide, inflammatory response, gamma-glutamylcyclotransferase activity, and genes coding components of cell-surface and external side of plasma membrane (Fig. 3B, 4B and 5B). Interestingly, PMNs from both old WT and CD73KO but not young WT mice showed down-regulation of NF-κβ signaling regulation upon *S. pneumoniae* challenge. In summary, the majority of down-regulated DEGs across all three PMN types belonged to the categories of transcription regulators and immune regulators.

### *S. pneumoniae* induces changes in lncRNA expression in the absence of CD73

Further analysis identified a total of 22 lncRNAs which were either significantly up- or down-regulated in CD73KO PMNs upon pneumococcal challenge (Fig. 7). We observed a lower number of lncRNAs (n=5) in WT PMNs from young mice (Fig 8), while PMNs from WT old mice had none. We made several searches in all available gene ontology (GO) and annotation databases and found to the best of our knowledge that these lncRNAs have not been previously functionally annotated. We therefore performed prediction and network analysis (see materials and methods) of CD73KO specific lncRNAs and found a total of 105 potential target interactions including 3 genes (*Il10, Icam1 and Rora*) identified in our RNA sequencing analysis (Fig 7 and Supplementary Table VIII). Since lncRNAs could directly bind to the target mRNA through complementary base pairing and thus determine the regulation of gene expression, we therefore inferred the biological functions of our lncRNAs based on their direct interaction with the gene targets, which, in turn, perturb the biological process in the disease pathway. For example, *Gm37747* can bind to several gene targets including *Cers6-205, Atp8a1-207, Spc25, Lpr2, Il10, Icam1, Atf3* and *Ldb2-204* which perturb signal transduction, regulation of cell adhesion and cellular response to tumor necrosis factor. This would signify that *Gm37747* is an important lncRNAs in these pathways. Importantly, we identified 5 pathways (Longevity regulation pathway, MAPK signaling pathway, Apoptosis signaling pathway, Nuclear receptor transcription pathway and Metabolic pathway) which were regulated by these lncRNAs (Fig.7). Among the predicted biological processes (Fig.7) were several signaling pathways including Signal transduction by protein phosphorylation cascade, Positive regulation of MAPK, Interferon signaling, Cytokine signaling in immune system, Cellular response to Tumor Necrosis Factor, Positive regulation of protein kinase C and Negative regulation of protein kinase B signaling. For young WT PMNs, lncRNA network analysis predicted different target genes (Supplementary Table VIII) but only one biological process was found and it connected to *Reep3* in the network (Olfactory signaling pathway and cellular component organization or biogenesis) (Fig. 8). These findings suggest that during *S. pneumoniae* infection, expression of lncRNAs in PMNs is controlled by CD73.

**Figure 7.**
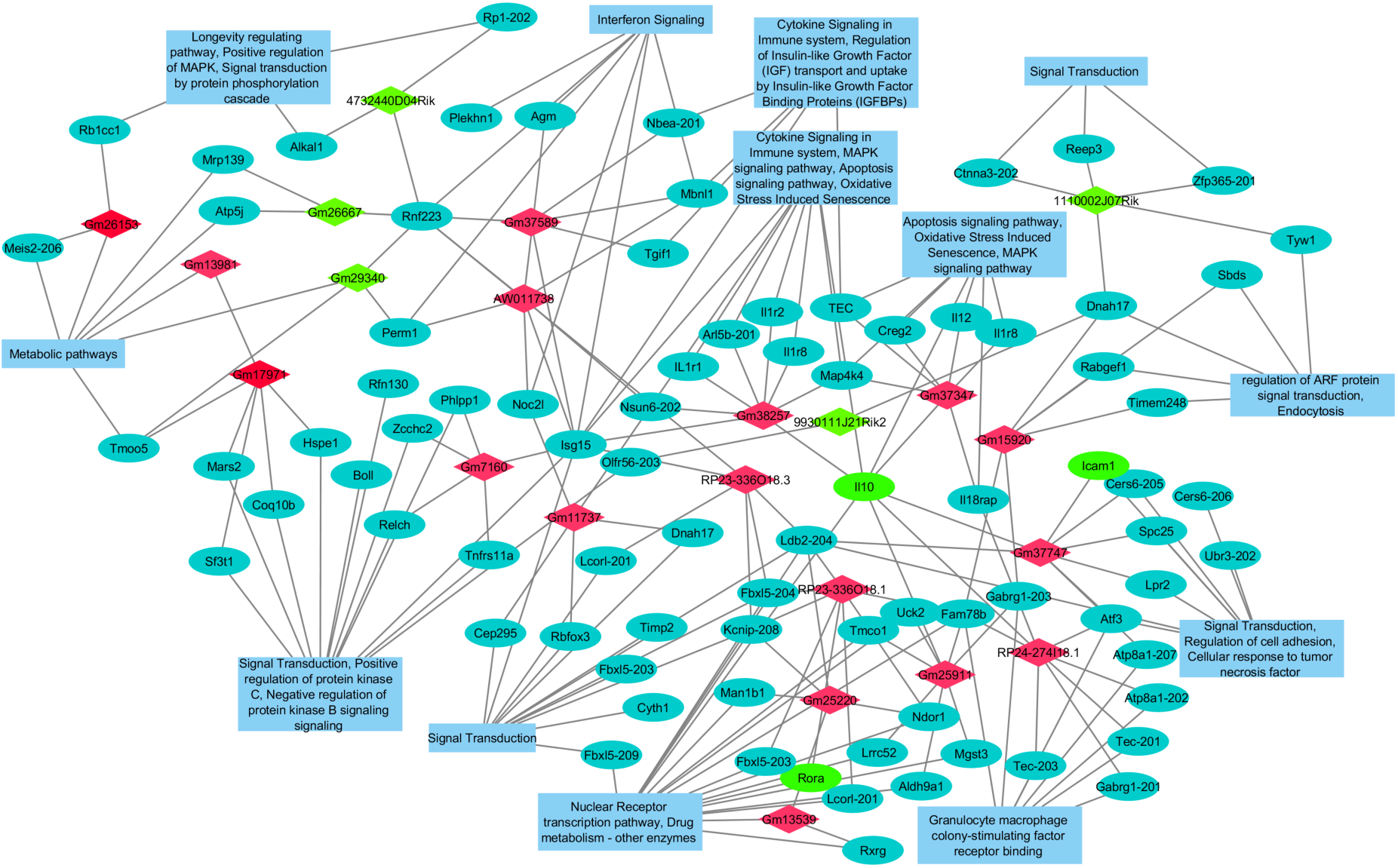
CD73KO PMN specific lncRNA-target network and biological process. In the network, red diamonds represent significantly up-regulated lncRNAs in CD73KO PMNs in response to *S. pneumoniae* infection, while green diamonds are the down-regulated lncRNAs. Blue ovals indicate predicted gene targets for the lncRNAs while the green ovals are the genes identified in actual RNA sequencing analysis. The big blue rectangles represent the predicted significantly impacted biological processes and pathways.

**Figure 8.**
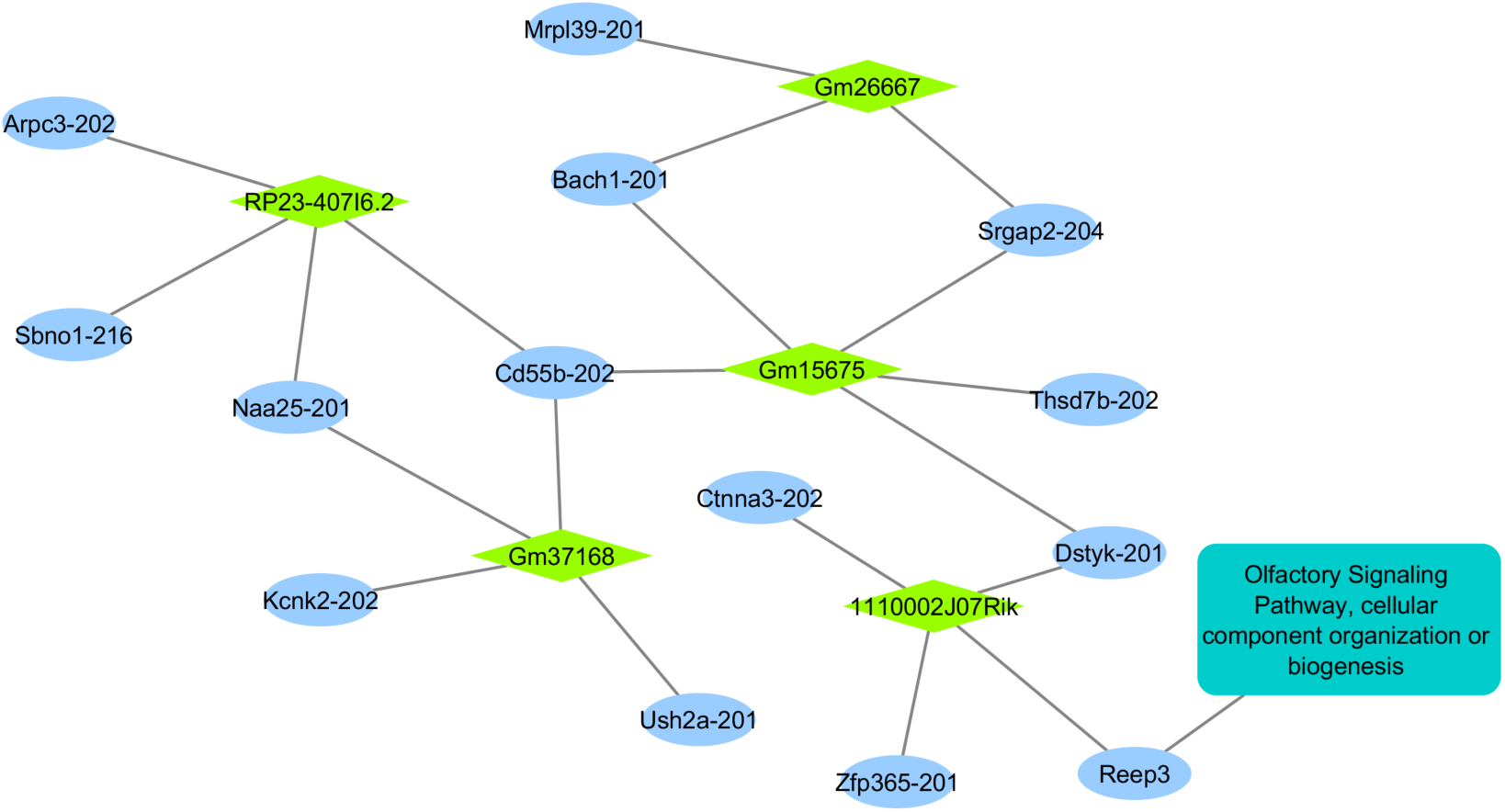
Young WT PMN specific lncRNA-target network and biological process. In the network, green diamonds represent the down-regulated lncRNAs in young WT PMNs in response to *S. pneumoniae* infection. Blue ovals are predicted gene targets for the lncRNAs. The big rectangle represents the predicted significantly impacted biological pathway.

### RT-PCR validation

To validate our RNA sequencing data, we tested the expression of a subset of differentially expressed genes through RT-PCR. The selection of genes tested was based on following categories: role in PMN function (*Il10* and *Adora2b*), role in MAPK pathways (*Fos*, *Jun*, *Hspa1a*, *Atf3*) or selected randomly (*Rrad* and *Rgl1*). The same samples on which RNA sequencing was performed were converted to cDNA for RT-PCR validation. Data were analyzed by the comparative threshold cycle (2 ^-ΔΔCT^) method, normalizing the CT values obtained for target gene expression to those for GAPDH of the same sample. Average of fold change values of target mRNA expression in infected samples was calculated relative to un-infected controls and then converted to log2 scale, as described for the RNA sequencing data. Multiple targets were tested in the CD73KO RNA samples as this group showed the strongest transcriptional response to *S. pneumoniae* challenge. Overall, the average log2 fold change values obtained during RT-PCR and RNA sequencing were consistent for the tested target genes (Fig. S2A, Fig. 9C and 9D) with a Pearson correlation coefficient of 0.8632 and *p*-value <0.01 (Fig. S2B).

**Figure 9.**
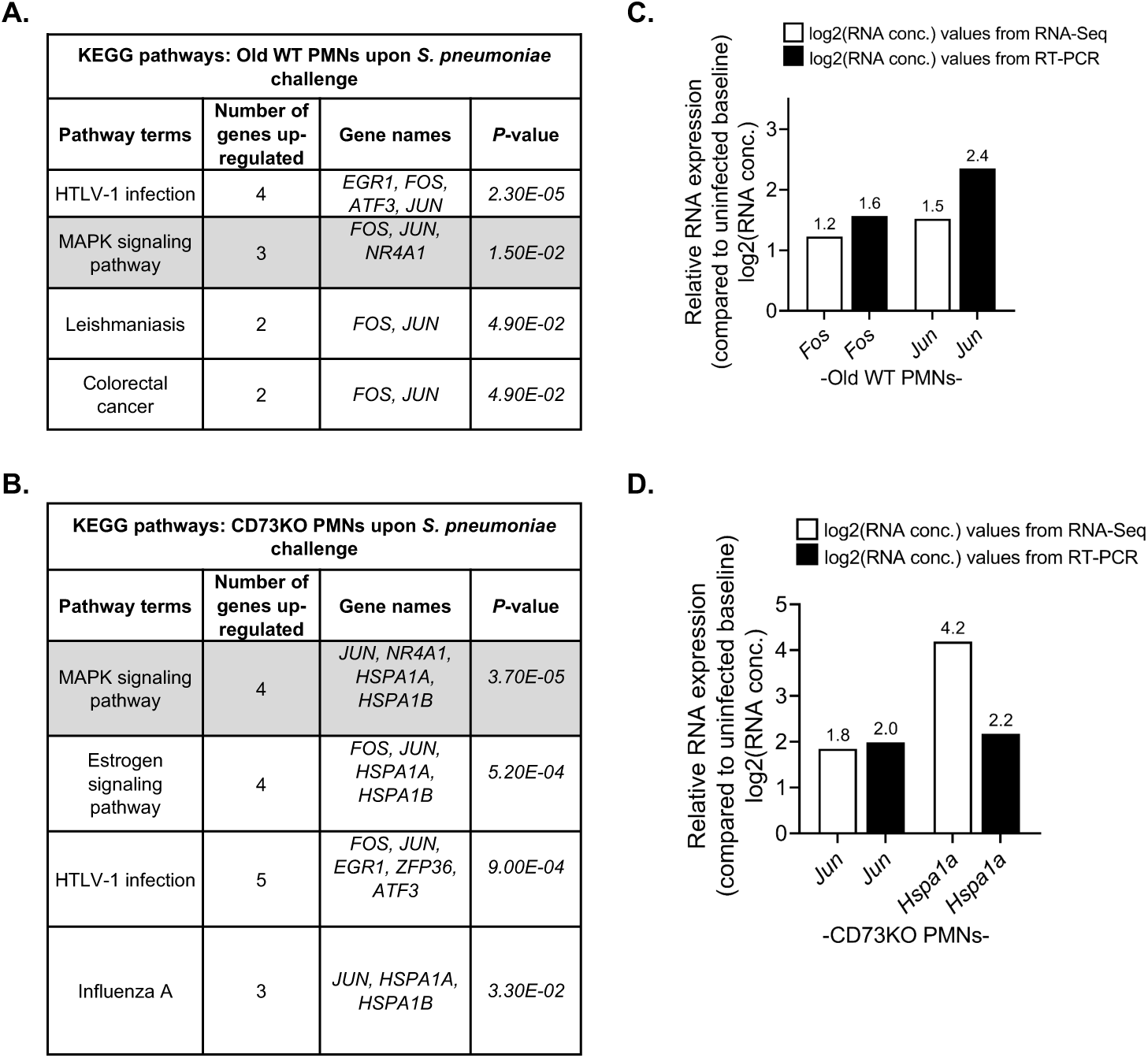
Validation of MAPK signaling pathway differentially expressed genes by real-time PCR. Lists of KEGG pathways and the corresponding genes retrieved from DAVID software using significantly up-regulated DEGs (log2FC ≥ 1.0, FDR < 0.05) in PMNs isolated from the bone marrow of old WT (A) or CD73KO (B) mice in response to infection with *S. pneumoniae* TIGR4 compared to mock-infected control are shown. Expression of select up-regulated DEGs corresponding to MAPK signaling pathway identified during RNA sequencing (white bars) was validated by RT-PCR (black bars) for old WT (C) and CD73KO PMNs (D). The data shown are the log2 value of the average of fold change values of target mRNA expression in infected samples relative to mock-infected controls. Relative fold change in target mRNA expression was calculated using three separate biological samples. Data were analyzed by the comparative threshold cycle (2 -ΔΔCT) method, normalizing the CT values obtained for target gene expression to those for GAPDH of the same sample.

### The MAPK signaling pathway is differentially up-regulated in PMNs from CD73KO and old mice in response to *S. pneumoniae* infection

DEGs significantly down-regulated or up-regulated in PMNs from young WT, old WT and CD73KO PMNs upon pneumococcal challenge were analyzed separately to identify pathways responsive to *S. pneumoniae* challenge. As the number of significant DEGs (FDR value < 0.05) identified was low, we first used a liberal approach to perform functional category analysis using DAVID where all functional categories and pathways with *p*-value < 0.05 were considered significant. We found that for PMNs from young WT mice, Autoimmune thyroid disease and Cytokine-cytokine receptor interaction pathway terms were down-regulated (Fig. 3B). No down-regulated KEGG pathway was observed in PMNs from old WT mice. In contrast, down-regulated pathways were identified in CD73KO PMNs (Table III) and included; Malaria, Cytokine-cytokine receptor interaction, Chemokine-signaling pathway, Allograft rejection, Cell adhesion molecules and Autoimmune thyroid disease (Fig. 5B). When comparing pathways that were up-regulated, we did not find any in PMNs from young WT mice. In contrast, in PMNs from old WT mice, the up-regulated pathway terms included HTLV-1 infection, MAPK signaling pathway, Leishmaniasis and Colorectal cancer (Fig. 4C), while the up-regulated pathways in CD73KO PMNs included MAPK signaling pathway, Estrogen signaling pathway, HTLV-1 infection and Influenza A (Fig. 5C).

**Table III.** Down-regulated KEGG pathways in PMNs isolated from young WT or CD73KO mice in response to *S. pneumoniae* infection.

| Downregulated KEGG pathways: Young WT PMNs upon *S. pneumoniae* challenge |  |  |  | Downregulated KEGG pathways: CD73KO PMNs upon *S. pneumoniae* challenge |  |  |
| --- | --- | --- | --- | --- | --- | --- |
| Pathway terms | Number of genes | Gene names | *P*-value | Number of genes | Gene names | *P*-value |
| Auto-immune thyroid disorder | 3 | *CTLA4*,<br>*GZMB*, *IL10* | *1.70E-02* | 3 | *GZMB*, *CD40*,<br>*IL10* | *3.1E-02* |
| Cytokine-cytokine receptor interaction | 4 | *CCR7*,<br>*CXCL16*,<br>*TNFRSF8*,<br>*IL10* | *3.0E-02* | 6 | *CCR7*, *CCL2*,<br>*CXCR5*,<br>*CXCL16*,<br>*CD40*, *IL10* | *2.20E-03* |
| Malaria |  |  |  | 4 | *ICAM1*,<br>*CCL2*, *CD40*,<br>*IL10* | *8.20E-04* |
| Chemokine signaling pathway |  |  |  | 5 | *ITK*, *CCR7*,<br>*CCL2*,<br>*CXCR5*,<br>*CXCL16* | *6.7E-03* |
| Allograft rejection |  |  |  | 3 | *GZMB*, *CD40*,<br>*IL10* | *2.0E-02* |
| Cell adhesion molecules (CAMs) |  |  |  | 4 | *PVR*, *ICAM1*,<br>*CD40*, *SDC4* | *2.5E-02* |

<sup>1</sup>
<sup>1</sup> Significantly ( $p \leq 0.05$ ) down-regulated KEGG pathways with genes involved and *p*-values based upon DAVID analysis of significantly down-regulated DEGs ( $\log_2\text{FC} \geq 1.0$ , $\text{FDR} < 0.05$ ) in PMNs isolated from young WT or CD73KO mice in response to *S. pneumoniae* TIGR4 infection compared to mock-challenged controls.

Importantly, PMNs from old WT and CD73KO mice shared two common up-regulated pathways including the MAPK signaling pathway (Fig. 9A and B). This pathway was also significantly up-regulated in CD73KO PMNs upon infection when the analysis was performed with FDR value < 0.05 criteria. KEGG analysis indicated that *S. pneumoniae* induced up-regulation of JNK as one of the common MAPK pathways in PMNs from old WT and CD73KO mice (Fig S3). We observed upregulation of *Fos* and *Jun* the components of activator protein-1 (AP-1) transcription complex which is regulated downstream of JNK signaling (37). Differential expression of select genes (*Fos, Jun,* and *Hspa1a*) in this pathway upon infection of PMNs from WT old and CD73KO mice was further confirmed using RT-PCR (Fig 9 C and D). To determine whether changes at the gene expression levels translated to functional differences in JNK pathway signaling, we quantified the proportion of c-Jun that undergoes phosphorylation in response to pneumococcal challenge. When phosphorylated, c-Jun forms part of the AP-1 transcription factor complex that is activated downstream of JNK signaling (37). We found increased phosphorylation of c-Jun in response to *S. pneumoniae* infection (Fig. 10) and importantly the portion of c-Jun that was phosphorylated was significantly higher in infected PMNs from old WT and CD73KO mice in comparison to young controls (Fig. 10B). These findings demonstrate age and CD73-driven changes in MAPK signaling in PMNs in response to pneumococcal infection.

**Figure 10.**
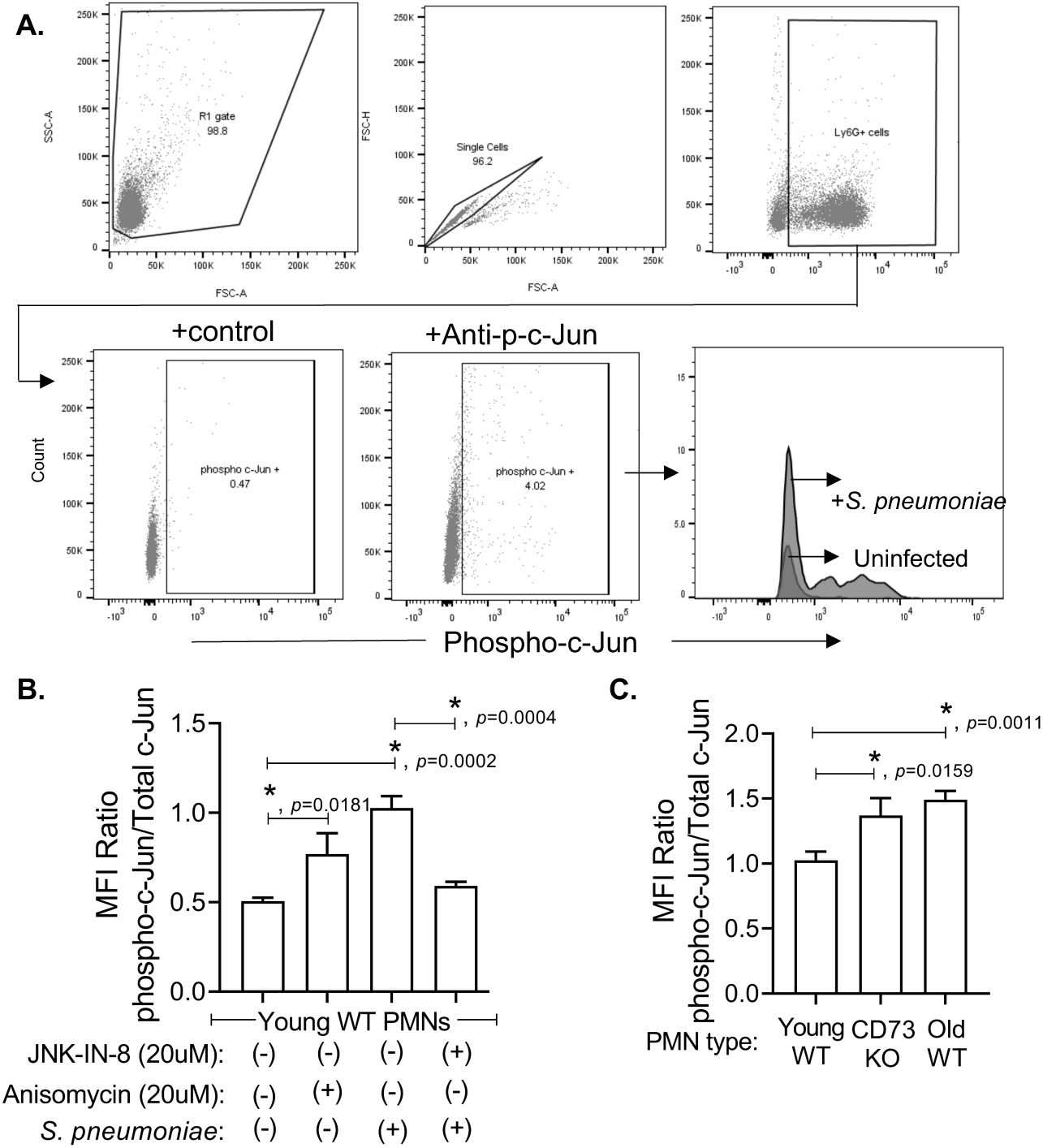
Phosphorylated c-Jun pools are higher in PMNs from old and CD73KO mice following *S. pneumoniae* infection. PMNs isolated from the bone marrow of the indicated strains of mice were incubated for 30 minutes at 37°C with *S. pneumoniae* TIGR4 pre-opsonized with matching sera at a MOI of 4 or mock-treated (uninfected) with 3% matching mouse sera only. Flow cytometry was used to determine the effect of bacterial infection on phospho-c-Jun (Ser73) levels. (A) The panel shows the gating strategy followed during analysis of flow cytometry in young WT mice. We gated on PMNs (Ly6G^+^ cells) and measured the expression (mean fluorescent intensity or MFI) of phospho-c-Jun (Ser73) and total-c-Jun. (B) PMNs from young WT mice were either mock-challenged, treated with Anisomycin (JNK/AP-1 pathway activator) or infected with *S. pneumoniae* in the absence or presence of JNK-IN-8 (JNK/AP-1 pathway inhibitor). The ratio of phosphorylated c-Jun with respect to the total cellular levels of c-Jun is presented. (C) PMNs from young WT, old WT and CD73KO mice were infected with *S. pneumoniae* and the ratio of phosphorylated c-Jun with respect to the total cellular levels of c-Jun was compared. Representative data (B and C) from one of five separate experiments where each condition was tested in triplicate (n=3 technical replicates) per experiment are shown. Asterisks indicate significant differences as calculated by Student’s t-test.

### Blocking JNK/AP-1 signaling pathway boosts bacterial killing in PMNs from old and CD73KO mice

We then wanted to explore whether the age and CD73-driven changes in the JNK MAPK pathway had an effect on PMN function. The JNK/AP-1 signaling pathway is well known for its role in stress-induced apoptotic cell death (38–40). Therefore, we tested whether there were differences in apoptosis between the mouse groups. Using Annexin-V-/Propidium iodide (PI) staining and flow cytometry we found that the percentage of apoptotic PMNs increased following infection (Fig S4A-C); however, there were no differences among the mouse groups. This was further confirmed using a lactate dehydrogenase (LDH) release assay (Fig S4D).

To determine whether JNK/AP-1 signaling played a role in PMN antibacterial function, we treated PMNs from young WT mice with the JNK stimulator Anisomycin and measured their ability to kill bacteria using our opsonophagocytic killing assay. The ability of Anisomycin to activate the JNK pathway was confirmed by measuring the extent of c-Jun phosphorylation by flow cytometry (Fig. 10B). Interestingly, we found a significant 2-fold reduction in the ability of PMNs from young mice to kill *S. pneumoniae* upon treatment with Anisomycin (Fig. 11A). As activation of JNK signaling blunted PMN antimicrobial function, we then asked whether the function of PMNs from old WT and CD73KO mice can be rescued by inhibiting this pathway. To do this, PMNs from old WT or CD73KO mice were treated with JNK-IN-8 or SR11302 prior to infection. JNK-IN-8 is a selective and high affinity inhibitor that irreversibly blocks the catalytic domain of JNK (41) while SR11302 is a selective inhibitor of AP-1 complex (42). The ability of JNK-IN-8 to inhibit phosphorylation of c-Jun was also confirmed by flow cytometry (Fig. 10B). We found that strikingly, treatment of PMNs with SR11302 or JNK-IN-8 significantly enhanced their ability to kill *S. pneumoniae* by 5- and 10-fold respectively, in both old and CD73KO mice (Fig 11 B and C). None of the JNK pathway inhibitors or activators had any significant effect on bacterial viability directly (Fig S5). These data indicate that blocking the JNK/AP-1 pathway reverses the defect in pneumococcal killing by PMNs from old WT and CD73KO mice.

**Figure 11.**
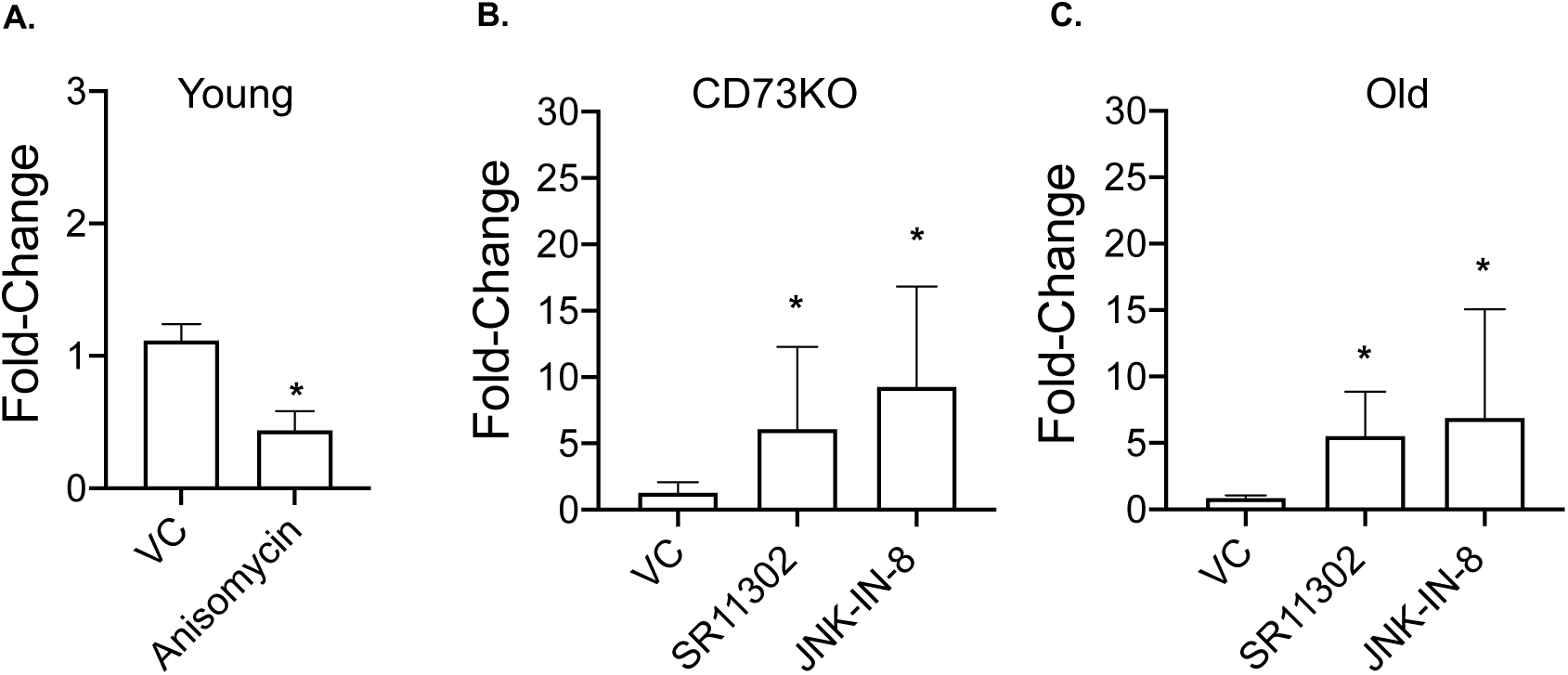
Blocking JNK/AP-1 pathway boosts the antimicrobial function of PMNs isolated from CD73KO and old WT mice. PMNs isolated from the bone marrow of young WT (A), CD73KO (B) and old WT (C) mice were treated with the indicated JNK-stimulator Anisomycin (20µM), JNK-inhibitor JNK-IN-8 (20µM), AP-1 inhibitor SR11302 (20µM) or HBSS+ (VC) for 30 minutes at 37°C. Treated PMNs were then challenged with *S. pneumoniae* TIGR4 strain pre-opsonized with homologous sera for 45 minutes at 37°C. Reactions were plated on blood agar plates and the percentage of bacteria killed compared to a no PMN control under the same condition was calculated. The fold-change in bacterial killing with respect to controls was then calculated by dividing the value of the treatment group by the vehicle control for each strain. Data shown are pooled from three separate experiments (n=3 biological replicates) where each condition was tested in triplicate (n=3 technical replicates) per experiment. Asterisks indicate significantly different from 1 by one-sample t-test.

## Discussion

PMN antimicrobial function declines with aging and is in part driven by changes in extracellular adenosine production and signaling (10). The aim of this study was to examine transcriptional changes in PMNs in response to *S. pneumoniae* infection across different hosts to better understand how aging impairs PMN function and what aspect of this was controlled by the extracellular adenosine-producing enzyme CD73. We found very limited differences in mRNA expression in mock-stimulated PMNs across the different hosts, indicating that either the intrinsic age-related defect in PMN function occurs at the protein level, or it is the transcriptional response following external stimulation which drives the difference in PMN responses, or both. In fact, *S. pneumoniae* infection triggered global transcriptional changes that were distinct across the different hosts.

A surprising finding was that there were 2-3-fold more down-regulated than up-regulated genes in response to infection across all host groups. Sixty percent of the up-regulated genes in WT mice were the same regardless of host age while the majority of the down-regulated DEGs were shared across the three different hosts suggesting an overall blunting of transcriptional activity and expression of only select transcripts in activated PMNs. This is reminiscent of stress responses observed in yeast cells where only genes required for resistance against a particular stressor are expressed while the rest are shut off, possibly to conserve energy (43–46). Overall, the number of DEGs in response to infection was not high (ranged from 45-103 genes across the different hosts), which is consistent with the lower amount of mRNA and overall transcriptional activity observed in PMNs as compared to other immune cells (47, 48). However, even these relatively moderate changes were key for efficient antimicrobial function as inhibition of transcription significantly impaired the ability of PMNs to kill pneumococci. It is possible that larger changes in gene expression would be observed with time as indicated by up-regulation of genes involved in transcription activation or transcription regulation across all hosts. Here, we limited our study to observing changes within forty minutes of infection due to concerns about the effects of bacterial infection on PMN viability in culture (49). In summary, this study shows that PMNs undergo transcriptional reprogramming which is required for their ability to efficiently kill bacteria.

CD73KO neutrophils displayed the strongest transcriptional response to *S. pneumoniae*, with 40% more differentially expressed genes during infection as compared to WT age-matched controls. This correlated with significant changes in expression of more than 20 lncRNAs in response to infection, 77% of which were up-regulated. In contrast, PMNs from young WT controls displayed only 5 differentially expressed lncRNAs, all of which were down-regulated, while PMNs from old mice had none. These findings suggest that during *S. pneumoniae* infection, lncRNA expression in PMNs is negatively controlled by CD73 or extracellular adenosine production. Extracellular adenosine was previously shown to activate expression of MEG3, a lncRNA in a liver cancer cell line (50). This study, to our knowledge, is the first to report a link between the EAD pathway and lncRNA expression in PMNs in response to infection. Furthermore, our data suggest that in the absence of CD73, changes in lncRNA expression dysregulates several biological processes in the cell, including those important for PMN antimicrobial activity. Recent studies have highlighted the role of lncRNAs in transcriptional regulation of inflammatory responses of several immune cells (51), including macrophages (52, 53) and human PMNs (54, 55). Interestingly, polymorphisms in LncRNAs expressed in neutrophils was associated with pneumococcal bacteremia in children in Kenya (56).

Of particular interest in our study, were genes that were up- and down-regulated only in PMNs from WT old and CD73KO mice but not in the young controls. Among genes that were down-regulated were *Rrad* and *CD40*, that have a role in oxidative responses. Binding of CD40 to its ligand activates downstream PI3K/NF-κβ leading to PMN oxidative burst (57) and defect in CD40 signaling is associated with blunted respiratory burst and antimicrobial activity in human PMNs (57). We previously found that CD73KO PMNs were defective in reactive oxygen species (ROS) production upon pneumococcal challenge (12). While PMNs from old mice do not show a defect in ROS production (10), aging is often associated with a buildup of reactive oxygen species, which if not controlled, can lead to cellular damage (58). Rrad (Ras-related associated with diabetes) is a GTP binding and calmodulin binding protein involved in reducing oxidative stress and preventing cellular senescence (59). Thus, reduction in Rrad expression could indicate an age-related decline in the ability of PMNs to counteract the oxidative stress induced following *S. pneumoniae* challenge. Among the genes up-regulated only upon infection in PMNs from old WT and CD73KO mice were *Slfn1* and *Nr4a1.* Slfn-1 is known for its role as inducer of cell cycle arrest in immune cells (60). Nr4a1 on the other hand belongs to family of nuclear receptor proteins that are rapidly induced under stress conditions and play an important role in DNA repair. Members of this family show aberrant expression in inflamed tissues and have emerged as key regulators of various diseases affecting the aging population (61). Interestingly, DNA damage and cell cycle arrest are characteristic features of cellular senescence (58). Overall, shared changes in gene expression in PMNs from old WT and CD73KO mice in response to infection, suggest an overall decline in the ability of these cells to aptly adapt to the infection-mediated stress, which in part is regulated by CD73.

KEGG pathway analysis showed that *S. pneumoniae* up-regulated MAPK-pathways in PMNs from both CD73KO and old mice but not in PMNs from young host. MAPK-pathways include JNK, p38, and ERK1/2, all of which regulate various cellular processes in response to external stimuli (62). Importantly, certain aspects of PMN function are attributed to different MAPK pathways. These include p38 MAPK and ERK mediated chemotaxis and respiratory burst (63, 64), MEK/ERK-mediated oxidative burst and phagocytosis (65) and p38 MAPK-mediated degranulation (66). Here, we observed upregulation of *Fos* and *Jun,* the components of activator protein-1 (AP-1) transcription complex which is regulated downstream of JNK signaling (37). We found that upon infection, c-Jun is phosphorylated in all mouse groups; however, the proportion of c-Jun undergoing phosphorylation was significantly higher in PMNs from old WT and CD73KO mice in comparison to young controls, indicating an increase in MAPK activation in these PMNs. Interestingly, host aging has been reported to be associated with an increase in basal levels of activation of other mitogen-activated protein kinase (MAPK) pathways including ERK1/2 and p38MAPK in PMNs (67–70). Importantly, these changes impair the ability of PMNs to respond to acute stimuli and impair their function (67, 71). For example, elevated activation of ERK1/2 impairs the ability of inflammatory signals to delay apoptosis in PMNs from elderly donors (68, 72). In fact, pharmacologically targeting these pathways has been shown to improve PMN function in elderly hosts. In sterile injury of the skin, oral administration of a p38 MAPK inhibitor resulted in enhanced PMN clearance in elderly donors (73). Similarly, here we found that stimulation of JNK/AP-1 blunted PMN anti-pneumococcal responses in young hosts while inhibition of this pathway rescued the ability of PMNs from old and CD73KO mice to kill *S. pneumoniae*, indicating that over-activation of JNK/AP-1 impairs PMN antimicrobial function. The role of JNK/AP-1 may be pathogen specific as inhibition of the JNK pathway decreased ROS production and release of NETs by PMNs in response to the Gram-negative bacteria *E. coli* and *P. aeruginosa* (74).

In conclusion, this study demonstrated the ability of PMNs to modify their gene expression to better adapt to bacterial infection and found that this capacity declines with age and is in part regulated by CD73. Importantly, we identified JNK/AP-1 signaling as a potential target for therapeutic intervention that can boost resistance of vulnerable hosts against *S. pneumoniae* infection.

## Acknowledgements

RNA sequencing was performed at Genomics and Bioinformatics Core facility at University at Buffalo. We thank Donald Yergeau for helpful discussions. We also thank Sujith A. Valiyaparambil for technical and logistical assistance.

## Notes

**Conflict of interest:** The authors have declared that no conflict of interest exists

1 This work supported by National Institute of Health grant R00AG051784 and a University at Buffalo Clinical and Translational Science Institute pilot award CTSA1153519 to ENBG. OBM was supported by an American Association of Immunologists Careers in Immunology fellowship and a College of Health Sciences and Technology bridge grant.

https://www.ncbi.nlm.nih.gov/geo/query/acc.cgi?acc=GSE150811

### Competing Interest Statement

The authors have declared no competing interest.

