## Supplemental materials for "Transcriptome profiling reveals CD73 and age-driven changes in neutrophil responses against *Streptococcus pneumoniae*"

**Supplemental Data**

**Supplemental Materials and Methods**

**Measurement of Apoptosis.** 2x$10^5$ PMNs were infected with pre-opsonized *S. pneumoniae* TIGR4 (MOI of 4) at 37° C for 5 and 40 minutes or mock treated with HBSS buffer containing 3% mouse sera. Flow cytometry was used to determine the percentage of live, apoptotic or necrotic cells using the FITC Annexin V apoptosis detection kit with PI (Biolegend) following the manufacturer's protocol.

**Lactate dehydrogenase (LDH) cell cytotoxicity assay.** Cytotoxic effect of *S. pneumoniae* infection on PMNs was evaluated by measuring the levels of the released cytoplasmic enzyme LDH. Briefly, 2x$10^5$ PMNs were infected with pre-opsonized *S. pneumoniae* TIGR4 strain (MOI of 4) at 37° C for 05 and 40 minutes or mock treated with HBSS buffer containing 3% mouse sera. The reactions were briefly centrifuged at 1000 rcf for 5 minutes and supernatants were collected to measure the release of LDH using the Cytotox 96 Assay kit (Promega, Madison, WI).

### Supplemental Figures and Legends

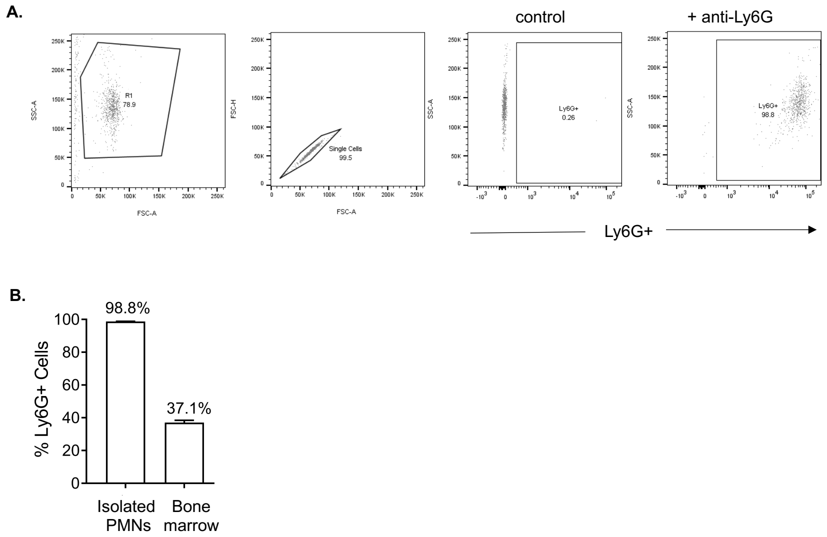


**Supplemental Figure 1. Negative selection yields a pure PMN population.** PMNs isolated from the bone marrow of young (WT) C57BL/6 mice were further enriched using a PMN negative selection kit (StemCell). Flow cytometry was used to determine the percentage of PMNs (Ly6G+ cells). (A) The panel shows the gating strategy followed during analysis. (B) The percentage of PMNs (gated on Ly6G+ cells) in the ultrapure PMN population compared to unfractionated bone marrow cells is shown.

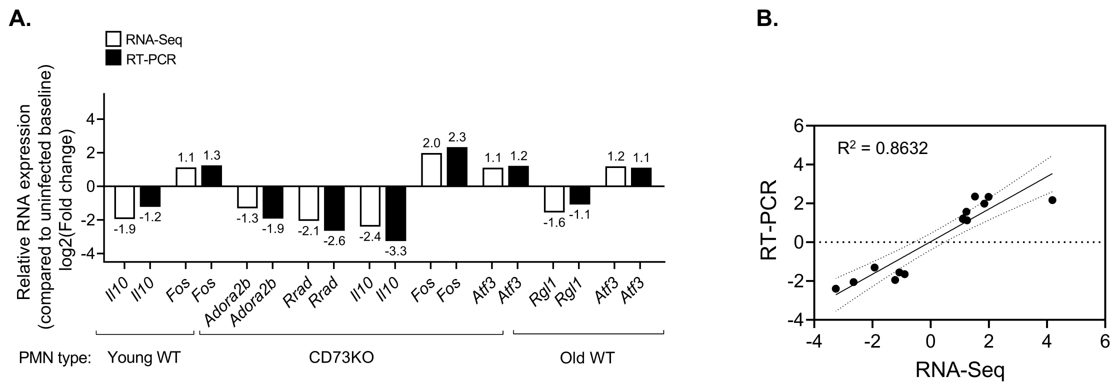


**Supplemental Figure 2. Validation of RNA sequencing data by real-time PCR.** (A) Expression of select up-regulated or down-regulated DEGs (indicated at the bottom of graph) identified during RNA sequencing was validated by RT-PCR. Data shown are the log2 value of the average of fold change values of target mRNA expression in infected samples relative to mock-infected controls. Relative fold change in target mRNA expression was calculated using three separate biological samples. Data were analyzed by the comparative threshold cycle ($2^{-\Delta\Delta CT}$) method, normalizing the CT values obtained for target gene expression to those for GAPDH of the same sample. (B) Correlation analysis between mean log2 (fold change) values obtained by RNA-Seq vs. qRT-PCR analysis was performed and Pearson correlation coefficient $R^2 = 0.8632$, *p*-value <0.01 was calculated.

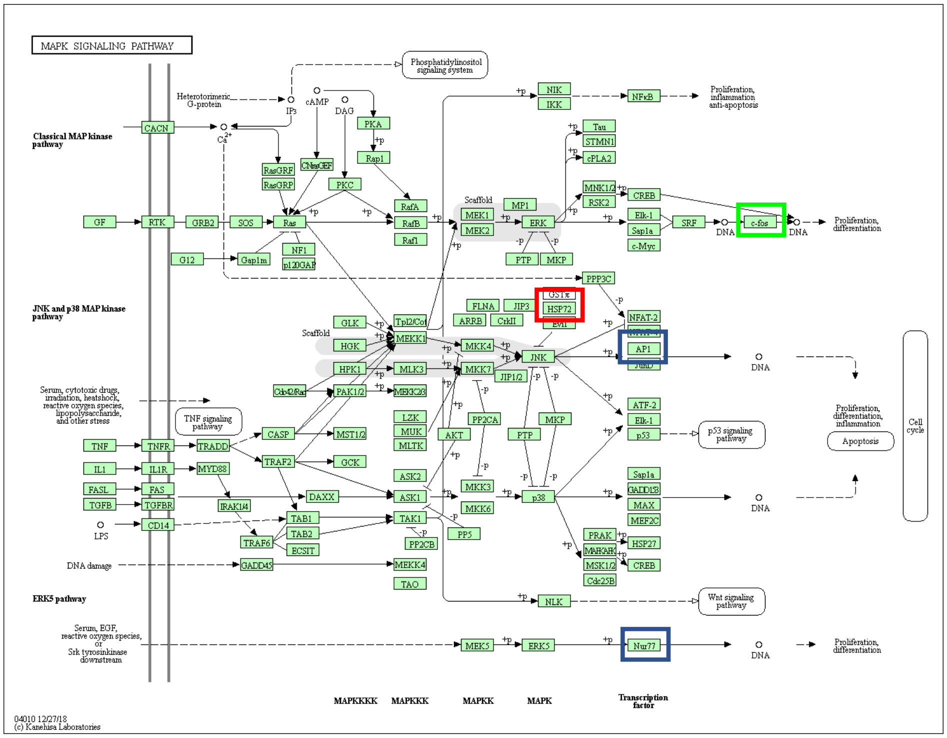


**Supplemental Figure 3. Differentially up-regulated MAPK pathway genes.** Diagram of MAPK pathway retrieved from the KEGG pathway analysis database and produced by DAVID using significantly up-regulated DEGs (log2FC ≥ 1.0, FDR < 0.05) from PMNs isolated from bone marrow of old WT or CD73KO mice in response to *S. pneumoniae* TIGR4 infection. Green boxes indicated up-regulated DEGs in old mice, salmon-colored boxes in CD73KO mice and blue boxes in both.

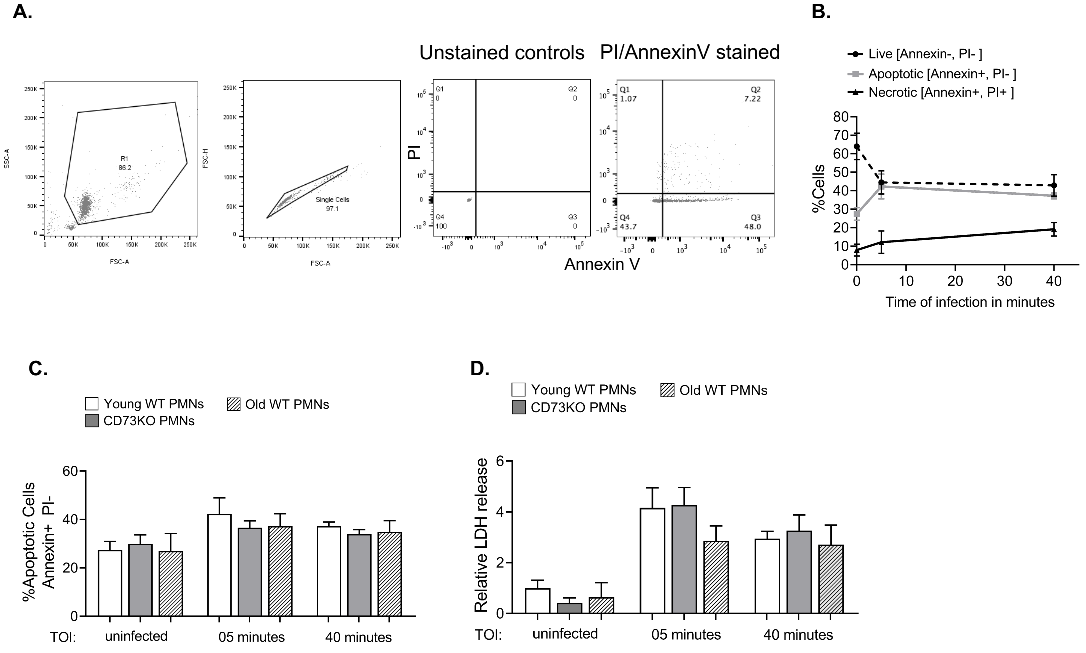


**Supplemental Figure 4. Aging and absence of CD73 have no impact on PMN viability in response to *S. pneumoniae* infection.** PMNs isolated from the bone marrow of young (WT) C57BL/6 mice were incubated for 5 and 40 minutes at 37°C with *S. pneumoniae* TIGR4 pre-opsonized with matching sera at a MOI of 2-4 or mock-treated (uninfected) with 3% matching mouse sera only. Flow cytometry was used to determine the effect of bacterial challenge on cell viability for the indicated time points. (A) The panel shows the gating strategy followed during flow cytometry analysis. The percentage of PMNs (gated on Ly6G+ cells) that were viable (PI- and Annexin V-), apoptotic (PI- and Annexin V +) or necrotic (PI+ and Annexin V+) are shown. (B) The percentage of live, apoptotic or necrotic PMNs isolated from young WT mice following *S. pneumoniae* infection were followed over the indicated timepoints. (C) The percentage of live, apoptotic or necrotic PMNs isolated from young WT, CD73KO and old WT mice (mock-infected and at 5 and 40 minutes post-infection were compared. (D) Infection-induced PMN cytotoxicity was quantified from the reaction supernatants by estimating the levels of LDH released at 5 and 40 minutes post-infection. Values of LDH released following challenge with *S. pneumoniae* were calculated relative to mock-infected controls. Data shown (B, C and D) were pooled from two separate experiments with one mouse per strain per condition tested in triplicate (n=3 technical replicates) per experiment.

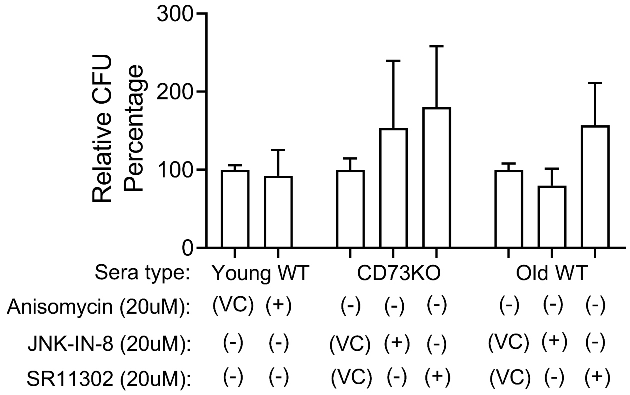


**Supplemental Figure 5. Effect of different drugs on bacterial viability.** *S. pneumoniae* TIGR4 were treated with HBSS+ (VC), Anisomycin (20μM), JNK-IN-8 (20μM) or SR11302 (20μM) for 45 minutes at 37°C. The number of viable bacteria was determined by plating on blood agar plates. Relative bacterial CFU were calculated against vehicle control and presented in terms of percentage increase or decrease in viability. Data shown are pooled from three separate experiments (n=3 biological replicates) where each condition was tested in triplicate (n=3 technical replicates) per experiment.

**Supplementary Table I.** RNA quality of the samples used in the study.

| Sample description | Conc. (ng/µl) | RIN |
|---|---|---|
| **Young WT (-S.p)** (Sample 1) | 64.77 | 8.5 |
| **Young WT (-S.p)** (Sample 2) | 68.94 | 7.7 |
| **Young WT (-S.p)** (Sample 3) | 107.31 | 10.0 |
| **Young WT (+S.p)** (Sample 4) | 103.95 | 9.3 |
| **Young WT (+S.p)** (Sample 5) | 73.11 | 9.8 |
| **Young WT (+S.p)** (Sample 6) | 192.35 | 9.4 |
| **Old WT (-S.p)** (Sample 7) | 65.19 | 8.1 |
| **Old WT (-S.p)** (Sample 8) | 61.09 | 9.2 |
| **Old WT (-S.p)** (Sample 9) | 72.21 | 9.2 |
| **Old WT (+S.p)** (Sample 10) | 58.52 | 6.9 |
| **Old WT (+S.p)** (Sample 11) | 75.19 | 9.4 |
| **Old WT (+S.p)** (Sample 12) | 93.81 | 8.8 |
| **CD73KO (-S.p)** (Sample 13) | 64.77 | 8.1 |
| **CD73KO (-S.p)** (Sample 14) | 58.10 | 9.7 |
| **CD73KO (+S.p)** (Sample 15) | 42.25 | 7.3 |
| **CD73KO (+S.p)** (Sample 16) | 75.6 | 6.8 |
| **CD73KO (+S.p)** (Sample 17) | 134.3 | 9.6 |

[1]

[1] -S.p indicates the mock-infected baseline controls; +S.p indicates the samples obtained from PMNs infected with *S. pneumoniae* TIGR4 strain.

**Supplementary Table II.** Biological Process, Cellular Component and Molecular Function terms differentially up-regulated in response to infection in PMNs from young WT mice.

| Biological Process (BP) terms | Genes involved in the process | Gene Count | P-Value |
|---|---|---|---|
| Negative regulation of transcription from RNA polymerase II promoter | *ZFP36, EGR1, PLK3, ATF3, JUN* | 5 | 1.60E-04 |
| Skeletal muscle cell differentiation | *EGR1, FOS, ATF3* | 3 | 2.50E-04 |
| Positive regulation of transcription from RNA polymerase II promoter | *EGR1, OSM, FOS, ATF3, JUN* | 5 | 5.30E-04 |
| Transcription from RNA polymerase II promoter | *EGR1, FOS, JUN* | 3 | 1.60E-03 |
| Positive regulation of transcription, DNA-templated | *EGR1, FOS, JUN, SERTAD3* | 4 | 1.60E-03 |
| Negative regulation of viral transcription | *ZFP36, TRIM21* | 2 | 5.30E-03 |
| Response to muscle stretch | *FOS, JUN* | 2 | 7.50E-03 |
| Positive regulation of pri-miRNA transcription from RNA polymerase II promoter | *FOS, JUN* | 2 | 9.70E-03 |
| Regulation of transcription, DNA-templated | *EGR1, FOS, ATF3, JUN, SERTAD3* | 5 | 1.20E-02 |
| Regulation of transcription from RNA polymerase II promoter | *ZFP36, FOS, ATF3* | 3 | 1.20E-02 |
| Response to radiation | *PLK3, JUN* | 2 | 1.30E-02 |
| Cellular response to hormone stimulus | *FOS, JUN* | 2 | 2.10E-02 |
| Positive regulation of cell proliferation | *OSM, ATF3, JUN* | 3 | 2.20E-02 |
| Response to cAMP | *FOS, JUN* | 2 | 2.20E-02 |
| Cellular response to calcium ion | *FOS, JUN* | 2 | 2.30E-02 |

| | | | |
|---|---|---|---|
| Positive regulation of neuron apoptotic process | *EGR1, JUN* | 2 | 3.00E-02 |
| Transforming growth factor beta receptor signaling pathway | *FOS, JUN* | 2 | 3.30E-02 |
| SMAD protein signal transduction | *FOS, JUN* | 2 | 3.40E-02 |
| Positive regulation of smooth muscle cell proliferation | *EGR1, JUN* | 2 | 3.50E-02 |
| Response to cytokine | *FOS, JUN* | 2 | 3.50E-02 |
| Transcription, DNA-templated | *EGR1, ATF3, JUN, SERTAD3* | 4 | 4.20E-02 |
| **Cellular Component (CC) terms** | **Genes involved in the process** | **Gene Count** | **P-Value** |
| Nucleus | *ZFP36, EGR1, FOS, PLK3, ATF3, JUN, SERTAD3, TRIM21* | 8 | 1.50E-03 |
| Cytoplasmic mRNA processing body | *ZFP36, TRIM21* | 2 | 3.00E-02 |
| Nucleolus | *PLK3, ATF3, SERTAD3* | 3 | 4.30E-02 |
| **Molecular Function (MF) terms** | **Genes involved in the process** | **Gene Count** | **P-Value** |
| Transcription regulatory region DNA binding | *EGR1, FOS, ATF3, JUN* | 4 | 1.30E-04 |
| DNA binding | *ZFP36, EGR1, FOS, ATF3, JUN, TRIM21* | 6 | 5.60E-04 |
| Double-stranded DNA binding | *EGR1, FOS, JUN* | 3 | 1.60E-03 |
| Sequence-specific DNA binding | *EGR1, FOS, ATF3, JUN* | 4 | 2.30E-03 |
| Transcription factor activity, sequence-specific DNA binding | *EGR1, FOS, ATF3, JUN* | 4 | 6.00E-03 |

| | | | |
|---|---|---|---|
| Transcriptional activator activity, RNA polymerase II core promoter proximal region sequence-specific binding | *EGR1, FOS, JUN* | 3 | 6.30E-03 |
| Transcription factor binding | *EGR1, FOS, JUN* | 3 | 9.90E-03 |
| RNA polymerase II core promoter proximal region sequence-specific DNA binding | *FOS, ATF3, JUN* | 3 | 1.10E-02 |
| R-SMAD binding | *FOS, JUN* | 2 | 1.10E-02 |
| Transcription factor activity, RNA polymerase II core promoter proximal region sequence-specific binding | *EGR1, JUN* | 2 | 1.10E-02 |
| Protein heterodimerization activity | *FOS, ATF3, JUN* | 3 | 2.20E-02 |
| Transcription regulatory region sequence-specific DNA binding | *EGR1, ATF3* | 2 | 2.40E-02 |
| RNA polymerase II core promoter sequence-specific DNA binding | *EGR1, FOS* | 2 | 3.00E-02 |
| Identical protein binding | *ATF3, JUN, TRIM21* | 3 | 3.10E-02 |

[1]

[1] List of Biological Process (BP), Cellular Component (CC) and Molecular Function (MF) terms ($p$ value ≤ 0.05) along with respective genes involved and gene count. The analysis is based upon list of significant DEGs which were up-regulated (log2FC ≥ 1.0, FDR < 0.05) in PMNs from the young mice challenged with *S. pneumoniae* strain TIGR4, compared to the mock-challenged control.

**Supplementary Table III.** Biological Process, Cellular Component and Molecular Function terms differentially down-regulated in response to infection in PMNs from young WT mice.

| Biological Process (BP) terms | Genes involved in the process | Gene Count | P-Value |
|---|---|---|---|
| Negative regulation of B cell proliferation | *CTLA4, PRDM1, IL10* | 3 | 8.60E-04 |
| Cellular response to lipopolysaccharide | *HAVCR2, ICAM1, CXCL16, IL10, TNIP3* | 5 | 1.60E-03 |
| Adaptive immune response | *HAVCR2, RNF125, CTLA4, PRDM1* | 4 | 4.70E-03 |
| Inflammatory response | *HAVCR2, CCR7, MEFV, TNFRSF8, IL10* | 5 | 9.60E-03 |
| T cell antigen processing and presentation | *ICAM1, TREML4* | 2 | 9.70E-03 |
| Negative regulation of myeloid dendritic cell activation | *HAVCR2, IL10* | 2 | 9.70E-03 |
| Immune system process | *HAVCR2, RNF125, MEFV, CTLA4, PRDM1* | 5 | 1.40E-02 |
| Cellular response to interleukin-1 | *ICAM1, SAA3, RORA* | 3 | 1.60E-02 |
| Regulation of cell proliferation | *SGK1, JAG2, TNFRSF8, PRDM1* | 4 | 1.80E-02 |
| Negative regulation of inflammatory response | *MEFV, RORA, IL10* | 3 | 1.90E-02 |
| Immune response | *CCR7, CTLA4, TNFRSF8, IL10* | 4 | 2.80E-02 |
| Circadian rhythm | *RORA, BHLHE40, ID3* | 3 | 2.80E-02 |
| Positive regulation of NF-kappaB transcription factor activity | *ICAM1, TRIM13, TNFRSF8* | 3 | 2.90E-02 |

| | | | |
|---|---|---|---|
| Cell adhesion mediated by integrin | *ICAM1, ITGA5* | 2 | 3.60E-02 |
| Inactivation of MAPK activity | *DUSP4, DUSP8* | 2 | 3.60E-02 |
| Negative regulation of interleukin-12 production | *MEFV, IL10* | 2 | 3.60E-02 |
| Notch signaling pathway | *CHAC1, JAG2, GZMB* | 3 | 3.70E-02 |
| Establishment of endothelial barrier | *ICAM1, PPP1R16B* | 2 | 3.80E-02 |
| Rhythmic process | *RORA, BHLHE40, ID3* | 3 | 3.90E-02 |
| Positive regulation of macrophage activation | *HAVCR2, IL10* | 2 | 4.30E-02 |
| Positive regulation of chemokine production | *HAVCR2, ADORA2B* | 2 | 4.80E-02 |
| Positive regulation of apoptotic process | *CTLA4, TNFRSF8, ID3, JMY* | 4 | 4.80E-02 |
| Positive regulation of GTPase activity | *ICAM1, CCR7, RGS1* | 3 | 4.90E-02 |
| **Cellular Component (CC) terms** | **Genes involved in the process** | **Gene Count** | **P-Value** |
| Intracellular | *RNF125, CCR7, MEFV, CHAC1, SAA3, TRIM13, SPATA13, CISH, JMY* | 9 | 2.10E-02 |
| External side of plasma membrane | *ICAM1, CCR7, ITGA5, CTLA4* | 4 | 3.20E-02 |
| Plasma membrane | *PVR, HAVCR2, ICAM1, SGK1, ADORA2B, ASTL, JAG2, CTLA4, SLCO2B1, OSCP1, CISH, CCR7, P2RY10, PPP1R16B, RGS1, ITGA5, SPATA13* | 17 | 4.80E-02 |
| Cell surface | *HAVCR2, ICAM1, CCR7, ADORA2B, ITGA5* | 5 | 4.80E-02 |

| Molecular Function (MF) terms | Genes involved in the process | Gene Count | P-Value |
|---|---|---|---|
| Gamma-glutamylcyclotransferase activity | *GGCT, CHAC1* | 2 | 9.40E-03 |
| MAP kinase tyrosine/serine/threonine phosphatase activity | *DUSP4, DUSP8* | 2 | 3.00E-02 |
| Notch binding | *CHAC1, JAG2* | 2 | 4.60E-02 |

[1]

[1] List of Biological Process (BP), Cellular Component (CC) and Molecular Function (MF) terms ($p$ value ≤ 0.05) along with respective genes involved and gene count. The analysis is based upon list of significant DEGs which were down-regulated (log2FC ≤ -1.0, FDR < 0.05) in PMNs from the young mice challenged with *S. pneumoniae* strain TIGR4, compared to the mock-challenged control.

**Supplementary Table IV.** Biological Process, Cellular Component and Molecular Function terms differentially up-regulated in response to infection in PMNs from old WT mice.

| Biological Process (BP) terms | Genes involved in the process | Gene Count | P-Value |
|---|---|---|---|
| Skeletal muscle cell differentiation | *EGR1, FOS, ATF3, NR4A1* | 4 | 1.50E-06 |
| Positive regulation of transcription from RNA polymerase II promoter | *EGR1, OSM, FOS, ATF3, JUN, NR4A1* | 6 | 2.40E-05 |
| Positive regulation of transcription, DNA-templated | *EGR1, FOS, JUN, SERTAD3, NR4A1* | 5 | 6.40E-05 |
| Negative regulation of transcription from RNA polymerase II promoter | *ZFP36, EGR1, ATF3, JUN, SLFN1* | 5 | 1.60E-04 |
| Regulation of transcription, DNA-templated | *EGR1, FOS, ATF3, JUN, SERTAD3, NR4A1* | 6 | 1.30E-03 |
| Transcription from RNA polymerase II promoter | *EGR1, FOS, JUN* | 3 | 1.60E-03 |
| Transcription, DNA-templated | *EGR1, ATF3, JUN, SERTAD3, NR4A1* | 5 | 5.80E-03 |
| Response to muscle stretch | *FOS, JUN* | 2 | 7.50E-03 |
| Positive regulation of pri-miRNA transcription from RNA polymerase II promoter | *FOS, JUN* | 2 | 9.70E-03 |
| Negative regulation of cell proliferation | *OSM, JUN, SLFN1* | 3 | 1.20E-02 |
| Regulation of transcription from RNA polymerase II promoter | *ZFP36, FOS, ATF3* | 3 | 1.20E-02 |
| Cellular response to fibroblast growth factor stimulus | *ZFP36, NR4A1* | 2 | 1.30E-02 |
| Cellular response to organic substance | *EGR1, NR4A1* | 2 | 1.50E-02 |
| Cellular response to hormone stimulus | *FOS, JUN* | 2 | 2.10E-02 |
| Positive regulation of cell proliferation | *OSM, ATF3, JUN* | 3 | 2.20E-02 |

| | | | |
|---|---|---|---|
| Response to cAMP | *FOS, JUN* | 2 | 2.20E-02 |
| Cellular response to calcium ion | *FOS, JUN* | 2 | 2.30E-02 |
| Positive regulation of endothelial cell proliferation | *JUN, NR4A1* | 2 | 2.90E-02 |
| Positive regulation of neuron apoptotic process | *EGR1, JUN* | 2 | 3.00E-02 |
| Transforming growth factor beta receptor signaling pathway | *FOS, JUN* | 2 | 3.30E-02 |
| SMAD protein signal transduction | *FOS, JUN* | 2 | 3.40E-02 |
| Positive regulation of smooth muscle cell proliferation | *EGR1, JUN* | 2 | 3.50E-02 |
| Response to cytokine | *FOS, JUN* | 2 | 3.50E-02 |
| **Cellular Component (CC) terms** | **Genes involved in the process** | **Gene Count** | **P-Value** |
| Nucleus | *ZFP36, EGR1, FOS, ATF3, JUN, SERTAD3, NR4A1, SLFN1* | 8 | 4.90E-03 |
| Transcription factor complex | *FOS, JUN, NR4A1* | 3 | 6.20E-03 |
| **Molecular Function (MF) terms** | **Genes involved in the process** | **Gene Count** | **P-Value** |
| Transcription regulatory region DNA binding | *EGR1, FOS, ATF3, JUN* | 4 | 1.90E-04 |
| Sequence-specific DNA binding | *EGR1, FOS, ATF3, JUN, NR4A1* | 5 | 1.90E-04 |
| Transcriptional activator activity, RNA polymerase II core promoter proximal region sequence-specific binding | *EGR1, FOS, JUN, NR4A1* | 4 | 2.90E-04 |
| Transcription factor binding | *EGR1, FOS, JUN, NR4A1* | 4 | 5.70E-04 |
| Transcription factor activity, sequence-specific DNA binding | *EGR1, FOS, ATF3, JUN, NR4A1* | 5 | 6.70E-04 |
| DNA binding | *EGR1, FOS, ATF3, JUN, NR4A1* | 6 | 1.20E-03 |
| Protein heterodimerization activity | *FOS, ATF3, JUN, NR4A1* | 4 | 1.90E-03 |
| Double-stranded DNA binding | *EGR1, FOS, JUN* | 3 | 2.10E-03 |
| Transcription factor activity, RNA polymerase II core promoter proximal region sequence-specific binding | *EGR1, JUN* | 2 | 1.20E-02 |

| | | | |
|---|---|---|---|
| R-SMAD binding | *FOS, JUN* | 2 | 1.20E-02 |
| RNA polymerase II core promoter proximal region sequence-specific DNA binding | *FOS, ATF3, JUN* | 3 | 1.40E-02 |
| Transcription regulatory region sequence-specific DNA binding | *EGR1, ATF3* | 2 | 2.70E-02 |
| RNA polymerase II core promoter sequence-specific DNA binding | *EGR1, FOS* | 2 | 3.40E-02 |

[1]

[1] List of Biological Process (BP), Cellular Component (CC) and Molecular Function (MF) terms ($p$ value $\leq 0.05$) along with respective genes involved and gene count. The analysis is based upon list of significant DEGs which were up-regulated (log2FC $\geq 1.0$, FDR $< 0.05$) in PMNs from the old mice challenged with *S. pneumoniae* strain TIGR4, compared to the mock-challenged control.

**Supplementary Table V.** Biological Process, Cellular Component and Molecular Function terms differentially down-regulated in response to infection in PMNs from old WT mice.

| **Biological Process (BP) terms** | **Genes involved in the process** | **Gene Count** | **P-Value** |
|---|---|---|---|
| Cellular response to lipopolysaccharide | *ICAM1, PLSCR2, CXCL16, CD40, TNIP3* | 5 | 5.50E-04 |
| Leukocyte cell-cell adhesion | *ICAM1, ITGA5, TNIP1* | 3 | 8.70E-04 |
| Negative regulation of I-kappaB kinase/NF-kappaB signaling | *RORA, TNIP1, TNIP3* | 3 | 2.30E-03 |
| T cell antigen processing and presentation | *ICAM1, TREML4* | 2 | 7.30E-03 |
| Positive regulation of NF-kappaB transcription factor activity | *ICAM1, TRIM13, CD40* | 3 | 1.70E-02 |
| Cell adhesion mediated by integrin | *ICAM1, ITGA5* | 2 | 2.70E-02 |
| Positive regulation of GTPase activity | *ICAM1, CCR7, CD40* | 3 | 2.90E-02 |
| Positive regulation of vascular endothelial growth factor production | *ADORA2B, RORA* | 2 | 4.30E-02 |
| Positive regulation of interleukin-12 production | *CCR7, CD40* | 2 | 4.60E-02 |
| Regulation of angiogenesis | *ADORA2B, ITGA5* | 2 | 5.00E-02 |
| **Cellular Component (CC) terms** | **Genes involved in the process** | **Gene Count** | **P-Value** |
| External side of plasma membrane | *ICAM1, CCR7, ITGA5, CD40* | 4 | 1.30E-02 |
| Cell surface | *ICAM1, CCR7, ADORA2B, ITGA5, CD40* | 5 | 1.60E-02 |
| Intracellular | *RNF125, CCR7, CHAC1, TRIM13, RRAD, RGL1, JMY* | 7 | 3.60E-02 |
| **Molecular Function (MF) terms** | **Genes involved in the process** | **Gene Count** | **P-Value** |

| | | | |
|---|---|---|---|
| Gamma-glutamylcyclotransferase activity | *GGCT, CHAC1* | 2 | 6.60E-03 |
| Polyubiquitin binding | *TNIP1, TNIP3* | 2 | 4.40E-02 |

[1]

[1] List of Biological Process (BP), Cellular Component (CC) and Molecular Function (MF) terms ($p$ value ≤ 0.05) along with respective genes involved and gene count. The analysis is based upon list of significant DEGs which were down-regulated (log2FC ≤ -1.0, FDR < 0.05) in PMNs from the old mice challenged with *S. pneumoniae* strain TIGR4, compared to the mock-challenged control.

**Supplementary Table VI.** Biological Process, Cellular Component and Molecular Function terms differentially up-regulated in response to infection in PMNs from CD73KO mice.

| Biological Process (BP) terms | Genes involved in the process | Gene Count | P-Value |
|---|---|---|---|
| Skeletal muscle cell differentiation | *EGR1, FOS, ATF3, BTG2, NR4A1* | 5 | 2.90E-07 |
| Negative regulation of transcription from RNA polymerase II promoter | *EGR1, ZFP36, PLK3, ATF3, BTG2, JUN, SNAI1, SLFN1* | 8 | 5.50E-06 |
| Negative regulation of cell proliferation | *OSM, BTG2, JUN, HSPA1A, SLFN1* | 5 | 6.00E-04 |
| Cellular response to hormone stimulus | *FOS, DUSP1, JUN* | 3 | 1.10E-03 |
| Response to cAMP | *FOS, DUSP1, JUN* | 3 | 1.30E-03 |
| Response to heat | *OSM, HSPA1A, HSPA1B* | 3 | 1.60E-03 |
| Positive regulation of endothelial cell proliferation | *JUN, F3, NR4A1* | 3 | 2.20E-03 |
| Negative regulation of apoptotic process | *PLK3, DUSP1, BTG2, JUN, HSPA1B* | 5 | 2.50E-03 |
| Positive regulation of transcription, DNA-templated | *EGR1, FOS, JUN, NR4A1, SNAI1* | 5 | 2.70E-03 |
| Positive regulation of transcription from RNA polymerase II promoter | *EGR1, OSM, FOS, ATF3, JUN, NR4A1* | 6 | 3.00E-03 |
| Positive regulation of apoptotic process | *DUSP1, JUN, RHOB, NR4A1* | 4 | 4.90E-03 |
| Regulation of transcription from RNA polymerase II promoter | *ZFP36, FOS, ATF3, SNAI1* | 4 | 7.80E-03 |
| Transcription from RNA polymerase II promoter | *EGR1, FOS, JUN* | 3 | 9.10E-03 |

| | | | |
|---|---|---|---|
| Positive regulation of nuclear-transcribed mRNA poly(A) tail shortening | *ZFP36, BTG2* | 2 | 1.30E-02 |
| Response to muscle stretch | *FOS, JUN* | 2 | 1.80E-02 |
| Positive regulation of cell proliferation | *OSM, ATF3, JUN, HLX* | 4 | 1.80E-02 |
| Positive regulation of pri-miRNA transcription from RNA polymerase II promoter | *FOS, JUN* | 2 | 2.30E-02 |
| Regulation of transcription, DNA-templated | *EGR1, FOS, ATF3, BTG2, JUN, HLX, NR4A1* | 7 | 2.50E-02 |
| Response to light stimulus | *FOS, DUSP1* | 2 | 2.60E-02 |
| mRNA catabolic process | *ZFP36, HSPA1A* | 2 | 2.90E-02 |
| Cellular response to fibroblast growth factor stimulus | *ZFP36, NR4A1* | 2 | 3.10E-02 |
| Response to radiation | *PLK3, JUN* | 2 | 3.10E-02 |
| Cellular response to organic substance | *EGR1, NR4A1* | 2 | 3.40E-02 |
| Negative regulation of cell cycle | *RHOB, NR4A1* | 2 | 3.70E-02 |
| Positive regulation of smooth muscle cell migration | *EGR1, F3* | 2 | 4.00E-02 |
| Transcription, DNA-templated | *EGR1, ATF3, BTG2, JUN, HLX, NR4A1* | 6 | 4.10E-02 |
| **Cellular Component (CC) terms** | **Genes involved in the process** | **Gene Count** | **P-Value** |
| Nucleus | *ZFP36, EGR1, NR4A1, HSPA1A, SNAI1, SLFN1, FOS, ATF3, PLK3, DUSP1,* | 14 | 7.60E-04 |

| | | | |
|---|---|---|---|
| | *TACSTD2, JUN, HLX, RHOB* | | |
| Transcription factor complex | *FOS, JUN, NR4A1* | 3 | 2.70E-02 |
| Cytoplasm | *EGR1, ZFP36, FOS, PLK3, DUSP1, F3, NR4A1, KLHL42, HSPA1A, SNAI1, SLFN1* | 11 | 7.00E-02 |
| Cytosol | *ZFP36, FOS, JUN, RHOB, HSPA1A* | 5 | 8.70E-02 |
| **Molecular Function (MF) terms** | **Genes involved in the process** | **Gene Count** | **P-Value** |
| Sequence-specific DNA binding | *EGR1, FOS, ATF3, JUN, HLX, NR4A1, SNAI1* | 7 | 4.00E-05 |
| Transcription regulatory region DNA binding | *EGR1, FOS, ATF3, JUN* | 4 | 1.90E-03 |
| DNA binding | *EGR1, ZFP36, FOS, ATF3, JUN, HLX, NR4A1, SNAI1* | 8 | 2.40E-03 |
| Transcriptional activator activity, RNA polymerase II core promoter proximal region sequence-specific binding | *EGR1, FOS, JUN, NR4A1* | 4 | 3.00E-03 |
| Transcription factor binding | *EGR1, FOS, JUN, NR4A1* | 4 | 5.70E-03 |
| Transcriptional repressor activity, RNA polymerase II core promoter proximal region sequence-specific binding | *ATF3, BTG2, SNAI1* | 3 | 8.50E-03 |
| Double-stranded DNA binding | *GR1, FOS, JUN* | 3 | 9.50E-03 |
| Transcription factor activity, sequence-specific DNA binding | *EGR1, FOS, ATF3, JUN, NR4A1* | 5 | 1.40E-02 |
| Protein heterodimerization activity | *FOS, ATF3, JUN, NR4A1* | 4 | 1.70E-02 |

| | | | |
|---|---|---|---|
| Protein binding | *ZFP36, PLK3, BTG2, JUN, RHOB, NR4A1, HSPA1A, HSPA1B, SNAI1, SLFN1* | 10 | 1.90E-02 |
| RNA polymerase II regulatory region sequence-specific DNA binding | *EGR1, ATF3, SNAI1* | 3 | 2.40E-02 |
| Transcription factor activity, RNA polymerase II core promoter proximal region sequence-specific binding | *EGR1, JUN* | 2 | 2.60E-02 |
| R-SMAD binding | *FOS, JUN* | 2 | 2.60E-02 |

[1]

[1] List of Biological Process (BP), Cellular Component (CC) and Molecular Function (MF) terms (*p* value ≤ 0.05) along with respective genes involved and gene count. The analysis is based upon list of significant DEGs which were up-regulated (log2FC ≥ 1.0, FDR < 0.05) in PMNs from the CD73KO mice challenged with *S. pneumoniae* strain TIGR4, compared to the mock-challenged control.

**Supplementary Table VII.** Biological Process, Cellular Component and Molecular Function terms differentially down-regulated in response to infection in PMNs from CD73KO mice.

| **Biological Process (BP) terms** | **Genes involved in the process** | **Gene Count** | **P-Value** |
|---|---|---|---|
| Cellular response to lipopolysaccharide | *HAVCR2, ICAM1, CCL2, CXCL16, CD40, IL10, TNIP3* | 7 | 5.40E-05 |
| Inflammatory response | *HAVCR2, CCR7, CCL2, GBP5, MEFV, CD40, TNIP1, IL10* | 8 | 1.10E-04 |
| Chemotaxis | *CCR7, CCL2, S1PR1, CXCR5, CXCL16* | 5 | 5.60E-04 |
| Positive regulation of GTPase activity | *ICAM1, CCR7, CCL2, S1PR1, CD40* | 5 | 1.20E-03 |
| Cellular response to interleukin-1 | *ICAM1, CCL2, SAA3, RORA* | 4 | 2.20E-03 |
| Leukocyte cell-cell adhesion | *ICAM1, ITGA5, TNIP1* | 3 | 2.70E-03 |
| Negative regulation of I-kappaB kinase/NF-kappaB signaling | *RORA, TNIP1, TNIP3* | 3 | 6.90E-03 |
| Immune system process | *HAVCR2, ITK, GBP5, MEFV, CD40, PRDM1* | 6 | 7.60E-03 |
| Regulation of cell adhesion | *ICAM1, S1PR1, NUAK1* | 3 | 8.00E-03 |
| T cell antigen processing and presentation | *ICAM1, TREML4* | 2 | 1.30E-02 |
| Negative regulation of myeloid dendritic cell activation | *HAVCR2, IL10* | 2 | 1.30E-02 |
| Positive regulation of cellular extravasation | *ICAM1, CCL2* | 2 | 1.90E-02 |
| Cell adhesion | *ICAM1, ICAM4, ASTL, ITGA5, NUAK1, MYBPC3* | 6 | 1.90E-02 |
| Positive regulation of endothelial cell proliferation | *PPP1R16B, CCL2, IL10* | 3 | 2.00E-02 |

| | | | |
|---|---|---|---|
| Negative regulation of inflammatory response | *MEFV, RORA, IL10* | 3 | 3.20E-02 |
| Cellular response to insulin stimulus | *SGK1, CCL2, TBC1D4* | 3 | 3.20E-02 |
| Innate immune response | *HAVCR2, ITK, MEFV, TRIM13, PRDM1* | 5 | 3.90E-02 |
| Single organismal cell-cell adhesion | *PVR, ICAM1, ICAM4* | 3 | 4.70E-02 |
| Cell adhesion mediated by integrin | *ICAM1, ITGA5* | 2 | 4.70E-02 |
| Negative regulation of interleukin-12 production | *MEFV, IL10* | 2 | 4.70E-02 |
| Positive regulation of NF-kappaB transcription factor activity | *ICAM1, TRIM13, CD40* | 3 | 4.80E-02 |
| Cellular response to tumor necrosis factor | *ICAM1, CCL2, RORA* | 3 | 4.90E-02 |
| Positive regulation of macrophage chemotaxis | *CCR7, CCL2* | 2 | 5.00E-02 |
| Establishment of endothelial barrier | *ICAM1, PPP1R16B* | 2 | 5.00E-02 |
| **Cellular Component (CC) terms** | **Genes involved in the process** | **Gene Count** | **P-Value** |
| External side of plasma membrane | *ICAM1, CCR7, S1PR1, CXCR5, ITGA5, CLEC2D, CD40* | 7 | 2.80E-04 |
| Cell surface | *HAVCR2, ICAM1, CCR7, ADORA2B, ITGA5, CD40, SDC4* | 7 | 8.80E-03 |

| | | | |
|---|---|---|---|
| Intracellular | *ITK, CCR7, MEFV, CHAC1, SAA3, TRIM13, TBC1D4, RRAD, SPATA13, RGL1, JMY* | 11 | 1.40E-02 |
| Plasma membrane | *HAVCR2, PVR, GPR84, ICAM1, SGK1, ADORA2B, ICAM4, ASTL, RRAD, CD40, SLCO2B1, CCR7, P2RY10, PPP1R16B, RND1, S1PR1, CXCR5, ITGA5, CLEC2D, SPATA13, EMP1* | 21 | 4.50E-02 |
| **Molecular Function (MF) terms** | **Genes involved in the process** | **Gene Count** | **P-Value** |
| Gamma-glutamylcyclotransferase activity | *GGCT, CHAC1* | 2 | 1.30E-02 |
| Signal transducer activity | *GPR84, CCR7, P2RY10, S1PR1, ADORA2B, CXCR5, TRIM13* | 7 | 1.70E-02 |
| Integrin binding | *ICAM1, ICAM4, ITGA5* | 3 | 4.10E-02 |
| Protein binding | *HAVCR2, PVR, SCHIP1, ITK, SGK1, GBP5, CCL2, ASTL, RRAD, GZMB, PIM2, RORA, CD40, JMY, ITGA5, BCL2A1A, CLEC2D, TREML4, ID3, PRDM1* | 20 | 4.90E-02 |

[1]

[1] List of Biological Process (BP), Cellular Component (CC) and Molecular Function (MF) terms (*p* value ≤ 0.05) along with respective genes involved and gene count. The analysis is based upon list of significant DEGs which were down-regulated (log2FC ≤ -1.0, FDR < 0.05) in PMNs from the CD73KO mice challenged with *S. pneumoniae* strain TIGR4, compared to the mock-challenged control.

**Supplementary Table VIII.** Predicted lncRNA-target interactions.

| lncRNA | Target | Normalized deltaG (ndG) | Minimum of local base-pairing interaction energies (kcal/mol) |
|---|---|---|---|
| Gm25220 | Ndor1 | -0.1716 | -41.3 |
| Gm25220 | Man1b1 | -0.0637 | -48.12 |
| Gm25220 | Gm13539 | -0.0465 | -53.06 |
| Gm25220 | Kcnip-208 | -0.2496 | -35.12 |
| Gm25220 | Ldb2-204 | -0.0179 | -41.1 |
| RP23-336O18.1 | Fbxl5-204 | -0.1079 | -47.02 |
| RP23-336O18.1 | Ndor1 | -0.2502 | -51.82 |
| RP23-336O18.1 | Fbxl5-203 | -0.1640 | -54.94 |
| RP23-336O18.1 | Rora | -0.2611 | -48.37 |
| RP23-336O18.1 | Ldb2-204 | -0.3765 | -53.77 |
| RP23-336O18.1 | Lcorl-201 | -0.3589 | -43.21 |
| RP23-336O18.1 | Kcnip-208 | -0.1653 | -47.93 |
| RP23-336O18.1 | Fam78b | -0.0972 | -34.45 |
| RP24-274I18.1 | Atp8a1-202 | -0.0423 | -56.09 |
| RP24-274I18.1 | Il18rap | -0.1753 | -40.6 |
| RP24-274I18.1 | Tec-203 | -0.1054 | -46.85 |
| RP24-274I18.1 | Gabrg1-201 | -0.0849 | -48.8 |
| RP24-274I18.1 | Atf3 | -0.0727 | -32.13 |
| RP24-274I18.1 | Il10 | -0.0692 | -49.22 |
| RP24-274I18.1 | Tec-201 | -0.0655 | -41.49 |
| RP24-274I18.1 | Fam78b | -0.4293 | -46.43 |
| Gm37747 | Cers6-205 | -0.4861 | -35.33 |
| Gm37747 | Atp8a1-207 | -0.2239 | -37.4 |
| Gm37747 | Spc25 | -0.0666 | -45.12 |
| Gm37747 | Lpr2 | -0.1450 | -42.34 |
| Gm37747 | Il10 | -0.0872 | -48.31 |
| Gm37747 | Icam1 | -0.3594 | -56.06 |
| Gm37747 | Atf3 | -0.0616 | -35.05 |
| Gm37747 | Ldb2-204 | -0.1465 | -39.91 |
| Gm25911 | Tmco1 | -0.1343 | -36.27 |
| Gm25911 | Uck2 | -0.1555 | -41.49 |
| Gm25911 | Fam78b | -0.1274 | -47.95 |
| Gm25911 | Aldh9a1 | -0.1353 | -53 |
| Gm25911 | Mgst3 | -0.1500 | -46.19 |
| Gm25911 | Lrrc52 | -0.1697 | -46.9 |

| | | | |
|---|---|---|---|
| Gm25911 | Gabrg1-203 | -0.1590 | -54.98 |
| Gm25911 | Il10 | -0.1890 | -56.35 |
| Gm38257 | Il1r2 | -0.1130 | -52.09 |
| Gm38257 | Map4k4 | -0.1100 | -41.9 |
| Gm38257 | IL1r1 | -0.1795 | -54.08 |
| Gm38257 | Il1r8 | -0.1826 | -46.01 |
| Gm38257 | Nsun6-202 | -0.1098 | -51.2 |
| Gm38257 | Arl5b-201 | -0.1066 | -43.35 |
| Gm38257 | Il10 | -0.1715 | -31.5 |
| Gm38257 | Isg15 | -0.1196 | -53.84 |
| Gm37347 | Il12 | -0.1478 | -47.34 |
| Gm37347 | TEC | -0.1039 | -49.79 |
| Gm37347 | Creg2 | -0.1001 | -38.36 |
| Gm37347 | Il18rap | -0.1920 | -35.83 |
| Gm37347 | Map4k4 | -0.1252 | -41.76 |
| Gm37347 | Il1r8 | -0.1224 | -43.08 |
| Gm37347 | Il10 | -0.1671 | -44.69 |
| AW011738 | Noc2l | -0.1964 | -31.7 |
| AW011738 | Nsun6-202 | -0.1128 | -42.75 |
| AW011738 | Agm | -0.3465 | -37.22 |
| AW011738 | Perm1 | -0.3647 | -36.51 |
| AW011738 | Isg15 | 0.1294 | -44.12 |
| AW011738 | Rnf223 | -0.2012 | -40.55 |
| AW011738 | Mbnl1 | -0.0416 | -39.82 |
| Gm37589 | Rnf223 | -0.0472 | -33.89 |
| Gm37589 | Mbnl1 | -0.0213 | -50.64 |
| Gm37589 | Tgif1 | -0.0105 | -52.73 |
| Gm37589 | Nbea-201 | -0.0875 | -40.53 |
| Gm37589 | Isg15 | -0.1924 | -44.22 |
| RP23-336O18.3 | Fbxl5-204 | -0.1602 | -32.08 |
| RP23-336O18.3 | Isg15 | -0.3761 | -41.99 |
| RP23-336O18.3 | Rnf223 | -0.1887 | -32.26 |
| RP23-336O18.3 | Ldb2-204 | -0.0932 | -29.89 |
| RP23-336O18.3 | Lcorl-201 | -0.0409 | -37.72 |
| RP23-336O18.3 | Kcnip-208 | -0.1987 | -35.86 |
| RP23-336O18.3 | Isg15 | -0.1082 | -48.27 |
| Gm11737 | Noc2l | -0.1415 | -53.63 |
| Gm11737 | Nsun6-202 | -0.0392 | -32.29 |
| Gm11737 | Cep295 | -0.0847 | -53.46 |

| | | | |
|---|---|---|---|
| Gm11737 | Rbfox3 | -0.0022 | -51.45 |
| Gm11737 | Dnah17 | -0.1566 | -40.02 |
| Gm15920 | Rabgef1 | -0.2408 | -52 |
| Gm15920 | Fam78b | -0.1127 | -54.82 |
| Gm15920 | Gabrg1-203 | -0.1125 | -52.53 |
| Gm15920 | Timem248 | -0.0434 | -55.66 |
| Gm15920 | Dnah17 | -0.1553 | -42.12 |
| Gm7160 | Zcchc2 | -0.2187 | -38.07 |
| Gm7160 | Phlpp1 | -0.1267 | -45.4 |
| Gm7160 | Tnfrs11a | -0.1241 | -31.1 |
| Gm7160 | Relch | -0.2767 | -29.59 |
| Gm7160 | Isg15 | -0.0959 | -34.05 |
| Gm17971 | Coq10b | -0.2056 | -39.23 |
| Gm17971 | Hspe1 | -0.4101 | -44.7 |
| Gm17971 | Mars2 | -0.0664 | -44.3 |
| Gm17971 | Tmoo5 | -0.0864 | -40.08 |
| Gm17971 | Gm13981 | -0.1652 | -36.78 |
| 9930111J21Rik2 | Olfr56-203 | -0.0909 | -44.31 |
| 9930111J21Rik2 | Dnah17 | 0.4949 | -41.94 |
| 4732440D04Rik | Rnf223 | 0.0622 | -38.57 |
| 4732440D04Rik | Alkal1 | -0.8466 | -41.07 |
| 4732440D04Rik | Rp1-202 | -0.0947 | -45.08 |
| 1110002J07Rik | Dnah17 | -0.0870 | -29.72 |
| 1110002J07Rik | Ctnna3-202 | -0.0800 | -42.99 |
| 1110002J07Rik | Zfp365-201 | -0.0461 | -32.76 |
| Gm29340 | Tmoo5 | -0.1432 | -37.21 |
| Gm29340 | Gm13981 | -0.0689 | -43.34 |
| Gm29340 | Rnf223 | -0.0947 | -54.94 |
| Gm26667 | Mrp139 | -0.0626 | -48.37 |
| Gm26667 | Rnf223 | -0.0702 | -53.77 |
| Gm26667 | Atp5j | -0.0286 | -42.39 |
| Gm15675 | Thsd7b-202 | -0.0392 | -51.62 |
| Gm15675 | Cd55b-202 | -0.0847 | -42.37 |
| Gm15675 | Srgap2-204 | -0.0022 | -50.31 |
| Gm15675 | Bach1-201 | -0.1566 | -44.23 |
| Gm15675 | Dstyk-201 | -0.2408 | -30.18 |
| RP23-407I6.2 | Arpc3-202 | -0.1127 | -43.92 |
| RP23-407I6.2 | Naa25-201 | -0.1125 | -32.68 |
| RP23-407I6.2 | Cd55b-202 | -0.0434 | -26.9 |

| | | | |
|---|---|---|---|
| RP23-407I6.2 | Sbno1-216 | -0.1010 | -32.71 |
| Gm37168 | Kcnk2-202 | -0.0664 | -35.63 |
| Gm37168 | Cd55b-202 | -0.0640 | -47.2 |
| Gm37168 | Naa25-201 | -0.1128 | -52.39 |
| Gm37168 | Ush2a-201 | -0.1745 | -36.09 |
| Gm26667 | Mrpl39-201 | -0.0599 | -53.21 |
| Gm26667 | Bach1-201 | -0.1056 | -50.41 |
| Gm26667 | Srgap2-204 | -0.1021 | -40.06 |

[1]

[1] Table showing the computational prediction of lncRNA-target interactions using LncTar software through free energy minimization. Using the normalized binding free energy (ndG), targets with value above -0.02 as cutoff were further used for network analysis.
